# Transcriptional and translational regulation of pathogenesis in Alzheimer’s disease model mice

**DOI:** 10.1101/2021.09.17.460831

**Authors:** Guillermo Eastman, Elizabeth R. Sharlow, John S. Lazo, George S. Bloom, José R. Sotelo-Silveira

## Abstract

**Background:** Defining the cellular mechanisms that drive Alzheimer’s disease (AD) pathogenesis and progression will be aided by studies defining how gene expression patterns change during pre-symptomatic AD and the ensuing periods of steadily declining cognition. Previous studies have emphasized changes in transcriptional regulation, but not translational regulation, leaving the ultimate results of gene expression alterations relatively unexplored in the context of AD.

**Objective:** To identify genes whose expression might be regulated at the transcriptional, and especially at the translational levels in AD, we analyzed gene expression in cerebral cortex of two AD model mouse strains, CVN (APPSwDI;NOS2^-/-^) and Tg2576 (APPSw), and their companion wild type (WT) strains at 6 months of age by tandem RNA-Seq and Ribo-Seq (ribosome profiling).

**Methods:** Identical starting pools of bulk RNA were used for RNA-Seq and Ribo-Seq. Differential gene expression analysis was performed at the transcriptional and translational levels separately, and also at the translational efficiency level. Regulated genes were functionally evaluated by gene ontology tools.

**Results:** Compared to WT mice, AD model mice had similar levels of transcriptional regulation, but displayed differences in translational regulation. A specific microglial signature associated with early stages of Aβ accumulation was up-regulated at both transcriptome and translatome levels in CVN mice. Although the two mice strains did not share many regulated genes, they showed common regulated pathways related to APP metabolism associated with neurotoxicity and neuroprotection.

**Conclusion:** This work represents the first genome-wide study of brain translational regulation in animal models of AD, and provides evidence of a tight and early translational regulation of gene expression controlling the balance between neuroprotective and neurodegenerative processes in brain.

## INTRODUCTION

Alzheimer’s disease (AD) is a progressive brain neurodegenerative disorder and the most common cause of dementia. Individuals with AD respond only marginally and briefly to currently available drugs and their long-term care is extremely costly. At the histopathological level, AD has well characterized hallmarks that include extracellular plaques made from amyloid-beta (Aβ) peptides, intraneuronal tangles made from the microtubule-associated protein, tau, synapse loss and neuron death [1].

A comprehensive understanding of the molecular mechanisms underlying AD is paramount for the development of novel therapies that can impede its onset and progression. In particular, impaired mRNA translation has been implicated in other neurological diseases [2–5], and there are reports linking Aβ and tau to dysregulated translation [6–12]. For example, local protein synthesis is altered in brain synaptosomes isolated from an AD mouse model that overproduces Aβ [6] and Aβ oligomers induce *de novo* synthesis of tau itself [8]. Also tau interacts with ribosomes *in vitro* [9] and *in vivo*, and decreases global translation [12] and the synthesis of ribosomal proteins [7]. Recently, an antisense transcript-mediated mechanism regulating tau translation has been described and implicated in human brain tau pathologies [13]. Furthermore, Aβ and tau work coordinately to regulate the mTORC1 complex [14, 15], which controls a plethora of cellular functions, including mRNA translation [16].

A diversity of genomics tools have been developed to understand neurodegenerative diseases [17], resulting in powerful studies to explore transcriptional changes associated with AD [18–21]. However, translational regulation has barely been examined, and other than one study about microglia [22], we are unaware of any work that directly compared transcriptional and translational regulation in the context of AD. Here, we describe the use of ribosome profiling, or Ribo-Seq [23–25], as a genomic screening technique to detect mRNA translation regulation in the brain cortex in transgenic 6 month old CVN (APPSwDI;NOS2^-/-^) and Tg2576 (APPSw) mice, both of which model human AD, and in wild type (WT) mice of the same strain backgrounds. At that age, CVN mice do not yet express any of the AD-like histological or behavioral phenotypes seen later in life ([CVN - Alzforum](https://www.alzforum.org/research-models/appswdi-x-nos2-knock-out)) but the Tg2576 strain already exhibits measurable cognitive impairment, reduced (long-term potentiation) LTP in the dentate gyrus and limited neuron loss (Tg2576 - Alzforum) Beginning with isolated, bulk cortical RNA, we performed RNA-Seq to define mRNA steady-state levels, and in tandem we produced and sequenced ribosome-associated mRNA footprints that revealed the exact positions of active ribosomes on mRNAs in the process of being translated into proteins. Quantitative analysis allowed us to estimate translational levels at a genome-wide level and to compare translational efficiency [24] between transgenic mice and their WT counterparts. We thereby uncovered translationally regulated genes with a complex signature that implicated genes involved in elevated Aβ production and neurotoxicity as well as in reducing the Aβ load in brain.

## MATERIALS AND METHODS

### Mice

Three 6 month old male mice from each of the following genotypes were used: CVN [26], Tg2576 [27] and their respective WT controls (C57/BL6 and B6;SJL, respectively; see Table 1). Animals were maintained, bred and euthanized in compliance with all policies of the Animal Care and Use Committee of the University of Virginia.

**Table 1.** AD model mice

| *Strain* | *Gene(s)* | *Mutations* | *Modification* | *WildTtype*<br>*Background*<br>*Strain* |
| --- | --- | --- | --- | --- |
| CVN<br>(APP<sub>SwDI</sub> X<br>NOS2<sup>-/-</sup>) | APP,<br>NOS2 | APP:<br>K670N/M671L,<br>E693Q, D694N | APP: Transgenic<br>NOS2: Knock-Out | C57/BL6 |
| Tg2576 | APP | APP: KM670/671NL | APP: Transgenic | B6;SJL |

\* APPSwDI x NOS2 Knock-out

### Transcriptomic and ribosome profiling of brain cortex

Cortices (∼250 mg each) were dissected from freshly removed brains in ice cold PBS containing 100 µg/ml cycloheximide (CHX; Sigma-Aldrich, catalog # 01810). The tissue was then cut into smaller pieces with a sterile scalpel and Dounce homogenized on ice in lysis buffer (5 mM Tris pH 7.5, 2.5 mM MgCl2, 1.5 mM KCl, 0.5% Triton X-100, 0.5% sodium deoxycholate, 2 mM 1,4-dithiothreitol and 100 µg/ml CHX), using 1 ml of lysis buffer per 100 mg of tissue. A transcriptome sample was then separated and total RNA was isolated using a *mirVana Total RNA Isolation Kit* (Invitrogen, catalog # AM1560) according to the vendor’s recommended protocol.

Ribo-Seq was performed as previously described [28, 29]. Briefly, lysate samples isolated as just described were centrifuged twice at 4°C at ∼17,000 g, for 1 and 10 minutes consecutively, to remove large cellular debris, nuclei, and mitochondria. The post-mitochondrial supernatant (∼1.6 ml; OD260 = 5-10 AU) was loaded onto a 12-33.5% (w/v) sucrose cushion prepared in polysome buffer (20 mM HEPES pH 7.5, 5 mM MgCl2, 100 mM KCl, 100 µg/ml CHX) and centrifuged for 2 hours at 36,000 rpm (222,228 gmax) in a Beckman SW41Ti rotor at 4°C. The polysome-enriched pellet was resuspended in polysome buffer and digested with 180-200 units of Benzonase nuclease (Millipore, catalog # E1014) for 10 minutes at room temperature to remove the RNA that was unprotected by ribosomes and thus produce protected ribosome footprints. Digestion was stopped by addition of 3 volumes of *mirVana Lysis Buffer*, and the RNA was isolated and then concentrated by overnight precipitation with 80% ethanol to maximize small RNA recovery. The concentrated RNA (10-15 µg per sample) was then size fractionated by electrophoresis using 15% polyacrylamide-urea gels (ThermoFisher Scientific, catalog # EC68852BOX) run at 200 volts in TBE (89 mM Tris-borate, pH 8.3, 2 mM EDTA) for 65 min to separate ribosome-protected mRNA footprints. Gels were stained using GelRed (Biotium, catalog # 41003) and circular agitation for 10 minutes in the dark. The ribosome footprint bands were identified using 26-mer and 34-mer RNA oligonucleotides [30], and excised in a dark room under UV light exposure. RNA recovery from gel slices was done overnight at room temperature by gentle mixing on a Nutator [30]. Size, quality and quantity of both transcriptome and translatome samples were evaluated in an Agilent 2100 Bioanalyzer using Nano and Small RNA kits (Agilent Technologies, catalog #s 5067-1511 and 5067-1548, respectively).

### Sequencing

All transcriptome and translatome samples were sequenced at BGI Tech Solutions (Hong Kong). Transcriptome samples were sequenced using an RNA-Seq quantification library protocol with ribosomal RNA (rRNA) removal library preparation, yielding at least 20 million paired-end (2x100 bp) reads. Translatome samples (ribosome footprints) were processed by small RNA library protocol, yielding at least 100 million single-end (50 bp) reads. Raw sequence data are available at the NCBI Sequence Read Archive (SRA; https://trace.ncbi.nlm.nih.gov/Traces/sra/) under BioProject ID PRJNA677972.

### Data Analysis

Quality control of sequence files was performed using *FastQC* [31] and then mapped against the *Mus musculus* genome (mm10/GRCm38 version) using *bowtie2* [32] and defaults parameters. Read counts over mRNAs or genes were estimated by *featureCounts* [33], and differential gene expression analysis of transcriptomes or translatomes was done separately using *edgeR* [34], comparing each strain with its respective WT parental strain. Normalized counts were exported, and translational efficiency was calculated and contrasted between AD model mouse strains and their WT counterparts using the *Xtail* R package [35]. For all comparisons (transcriptome, translatome and translational efficiency) differentially expressed genes (DEGs) were defined by a false discovery rate (FDR) adjusted p-value <0.05 and a fold change of >1 or >1.5, as indicated in each case. Functional interpretation and ontology enrichment analysis of DEG lists were performed using *Ingenuity Pathway Analysis* (IPA; QIAGEN Inc.) [36], online tools like *STRING* [37] and *g:Profiler* [38], and in-house software (manuscript in preparation; https://github.com/sradiouy/IdMiner). AmiGO2 database [39] was used to retrieve genes related to APP, Aβ and tau. Plots were generated using R, by general or specific packages, such as *pheatmap* (https://CRAN.R-project.org/package=pheatmap) and GOplot [40].

## RESULTS

### High quality datasets of brain transcriptome and translatome differentiate CVN and Tg2576 models

CVN and Tg2576 AD model mice, and the parental WT strains from which they were derived (Table 1 and Fig. 1A) were used to explore transcriptional and translational gene expression regulation in the brain cortex of six month old animals using RNA-Seq and Ribo-Seq, respectively (Fig. 1B). More than 20 million paired-end reads were obtained for transcriptomes and an average of 120 million reads were acquired for translatomes (Table S1), derived from total RNA and isolated ribosome footprints (Fig. S1), respectively. In the transcriptome samples, 88% of the reads aligned over the reference genome, of which 90% mapped to gene regions and 77% to mRNA regions (Table S1). On the other hand, in the translatome samples we mapped more than 10 million reads over mRNAs (Table S1). In this case, ribosomal RNA depletion was avoided to minimize protocol biases.

**Fig. 1.**
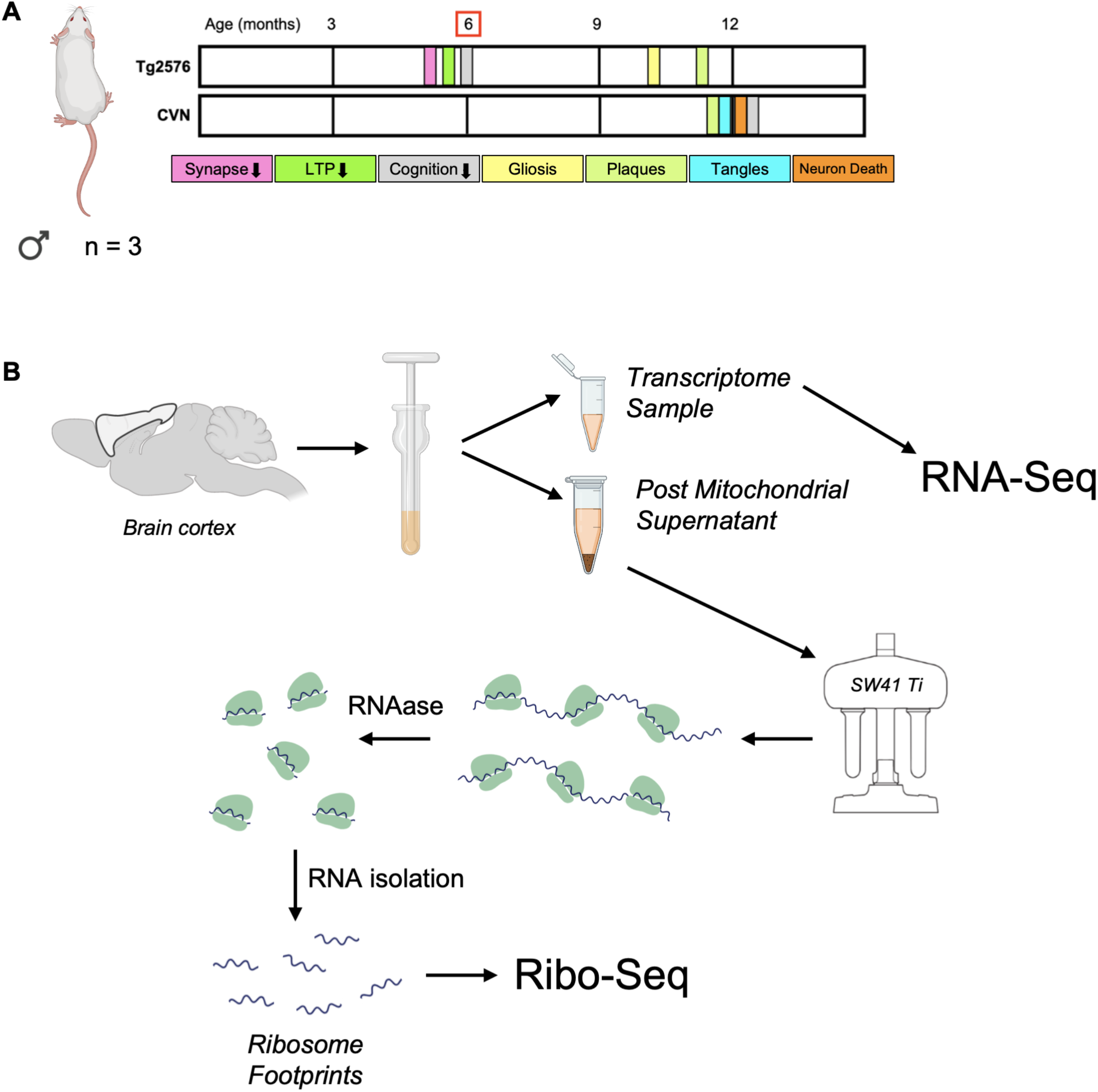
Protocol summary. A) Phenotypes of the CVN (adapted from <u>CVN - Alzforum</u>) and Tg2576 (adapted from <u>Tg2576 - Alzforum</u>) mice used in this study. Three 6 month old male mice were used for each transgenic strain, and for each corresponding WT strain (C57/Bl6 for CVN and B6;SJL for Tg2576). B) Sample preparation. Brain cortex was dissected and homogenized using a glass Dounce homogenizer to yield a transcriptome sample containing total RNA to be used for RNA-Seq, and a post-mitochondrial supernatant, which was ultracentrifuged to isolate polysomes. Ribosome footprints for Ribo-seq were isolated from the polysomes by an RNA protection assay that digested all RNA not encased inside the ribosomes. Further details are described in the Materials and Methods section.

Expression levels (CPM, counts per million) were estimated for more than 14 thousand different mRNAs in each sample above low/noise signal (Fig. S2). Inter-replicate correlations indicated high similarity within either transcriptome or translatome samples, but as expected, lower correlations were detected between RNA-Seq (transcriptome) and Ribo-Seq (translatome) data (Fig. S3). Quality control comparisons between transcriptome and translatome samples revealed the expected difference at the level of triplet periodicity and read distribution among mRNA features (Fig. S4 and S5). In addition, principal component analysis (PCA) showed a clear separation between genotypes for both RNA-Seq and Ribo-Seq datasets (Fig. S6).

### Gene expression is regulated transcriptionally and translationally in cortices of CVN and Tg2576 mice

We used the *edgeR* R package [34] to detect differences in cortical gene expression (p-adjusted value <0.05) with a directionally independent fold change (FC) of >1 between the transgenic mice and their respective WT controls at both the transcriptional and translational levels. For CVN versus their WT parental strain (C57/Bl6), we found 469 differentially expressed genes (DEGs) at the transcriptome level and 1165 DEGs at the translatome level. Of those, 240 and 377 genes had >1.5-fold differential expression by RNA-Seq and Ribo-Seq, respectively (Table 2 and Fig. 2). For Tg2576 versus their WT counterparts (B6;SJL), we found 343 DEGs at the transcriptome level and 135 at the translatome level with FC >1. Out of those totals, 140 transcriptionally regulated and 94 translationally regulated genes had a FC >1.5 (Table 2 and Fig. 3). Complete lists of the DEGs detected for CVN and Tg2576 mice are found in Tables S2 and S3, respectively.

**Fig. 2.**
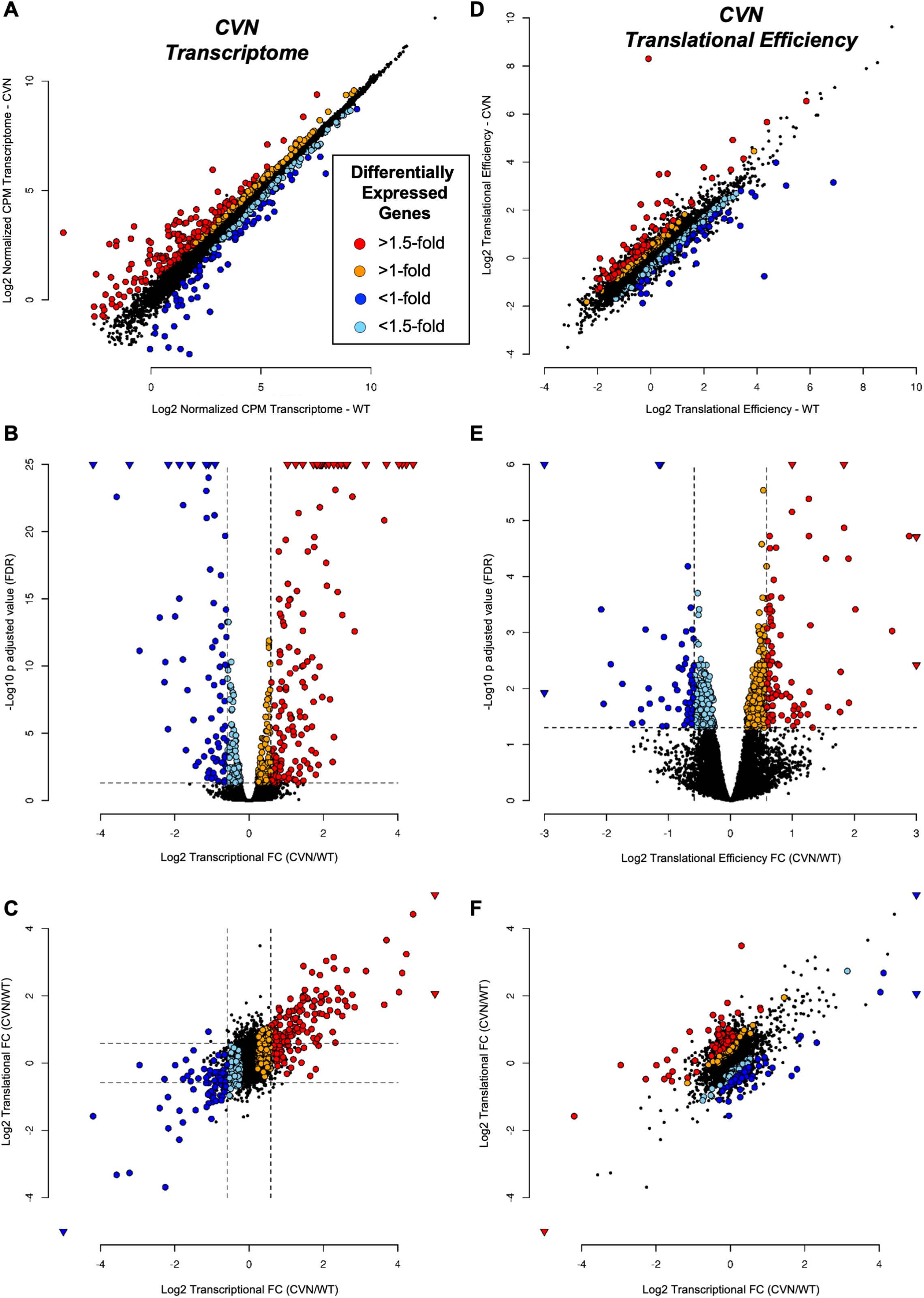
Differential expression in CVN versus WT (C57/Bl6) cortices determined by *edgeR* for transcriptomes (A-C), and by *Xtail* for translational efficiency (D-F). Scatter plots comparing normalized CPM expression between genotypes for transcriptome and translational efficiency values, respectively, are shown in (A) and (D). Volcano plots showing the relationship between fold change and p-adjusted values are shown in (B) and (E). Scatter plots comparing translational versus transcriptional fold change are shown in (C) and (F).

**Fig. 3.**
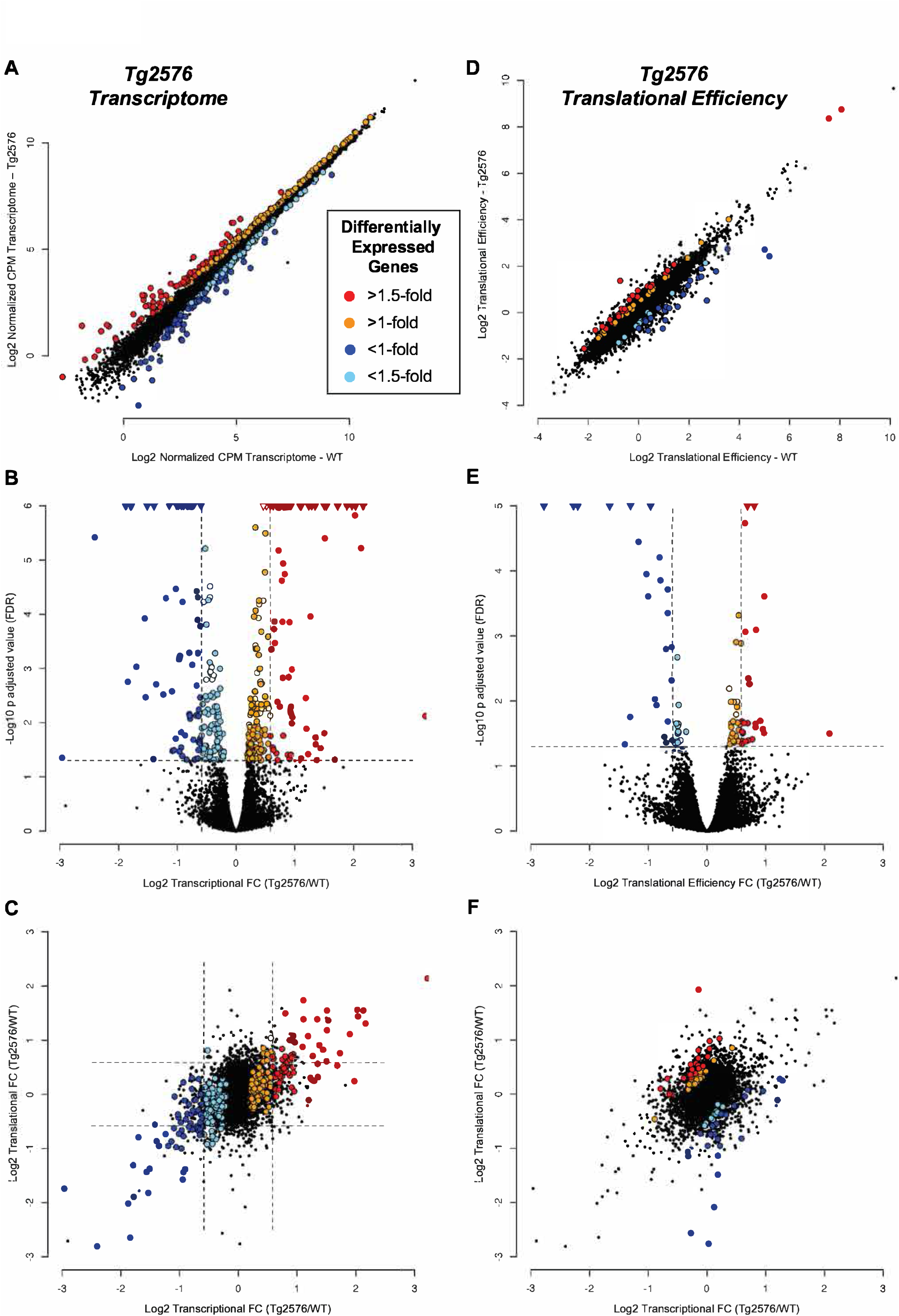
Differential expression in Tg2576 versus WT (B6;SJL) cortices determined by *edgeR* for transcriptomes (A-C), and by *Xtail* for translational efficiency (D-F). Scatter plots comparing normalized CPM expression between genotypes for transcriptome and translational efficiency values, respectively, are shown in (A) and (D). Volcano plots showing the relationship between fold change and p-adjusted values are shown in (B) and (E). Scatter plots comparing translational versus transcriptional fold change are shown in (C) and (F).

**Table 2.** Differentially expressed genes for CVN and Tg2576 mice versus their wild type background strain counterparts. Differentially expressed genes, defined by p-adjusted value <0.05, are separated by fold change (FC) intervals.

| **CVN vs WT** |  | *p-adjusted value < 0.05* |  |  |  |
| --- | --- | --- | --- | --- | --- |
|  |  | *FC > 1.5* | *FC > 1* | *FC < -1.5* | *FC < -1* |
| *edgeR* | RNA-Seq | 160 | 282 | 80 | 187 |
|  | Ribo-Seq | 300 | 830 | 77 | 335 |
| *Xtail* | Translational Efficiency | 88 | 491 | 56 | 306 |
| **Tg2576 vs WT** |  | *p-adjusted value < 0.05* |  |  |  |
|  |  | *FC > 1.5* | *FC > 1* | *FC < -1.5* | *FC < -1* |
| *edgeR* | RNA-Seq | 76 | 194 | 64 | 149 |
|  | Ribo-Seq | 46 | 66 | 48 | 69 |
| *Xtail* | Translational Efficiency | 25 | 50 | 25 | 37 |

To disentangle transcriptome from translatome regulation, we estimated translational efficiencies based on the ratios between translatome- and transcriptome-derived expressions levels for each transgenic strain and its WT parental counterpart. For this we used the *Xtail* R package [35], which revealed that CVN cortices contained 797 translationally regulated genes with FC >1, of which 144 had a FC >1.5 (Table 2 and Fig. 2). Similarly, Tg2576 cortices were found to contain 87 translationally regulated genes with FC >1, of which 50 had a FC >1.5 (Table 2 and Fig. 3).

The global distribution of DEGs values, the relationship between FC and p-adjusted values, and the association between transcriptome and translatome FCs are shown for both AD mouse models in Figures 2, 3, S7 and S8. As expected, DEGs were distributed along all expression levels (Fig. 2A and D, 3A and D) and the relationship between FC and p-adjusted values showed classic Volcano plots for both transcriptome and translational efficiency levels (Fig. 2B and E, 3B and E). When the FC at the translatome level was plotted against the transcriptome, a clear correlation was observed (Fig. 2C and 3C). For example, in CVN mice, 106 out of the 160 DEGs with a FC >1.5 at the transcriptome level were also DEGs at the same cutoff at the translatome level (Fig. S9). A similar correlation was evident for the transcriptionally down- regulated genes; 32 out of 80 were also down-regulated at the translational level (Fig. S9). Analogous results were also observed in Tg2576 mice (Fig. S9), and indicated the expected association between transcriptome and translatome samples.

By comparison, translationally regulated genes, although less in number than those regulated transcriptionally, showed a different pattern of regulation (Fig. 2F and 3F). In general, translational efficiency regulation involved genes with minimal changes at transcriptome levels (FC ∼1). This pattern was also apparent in the DEGs expression heatmap (Fig. 4). Transcriptome regulated genes expression levels showed classical patterns of regulation in contrast to the translational efficiency regulation. For example, similar expression levels for the transcriptome coupled with an increase or decrease for the translatome implied translational efficiency regulation. This scenario, among others, could be observed in the heatmaps of Figure 4. Heatmaps of regulated genes exclusively at the translational level are described in Figures S7D and S8D for CVN and Tg2576 mice, respectively. Collectively, these results demonstrate differential transcriptional and translational regulation of gene expression in both murine AD models.

**Fig. 4.**
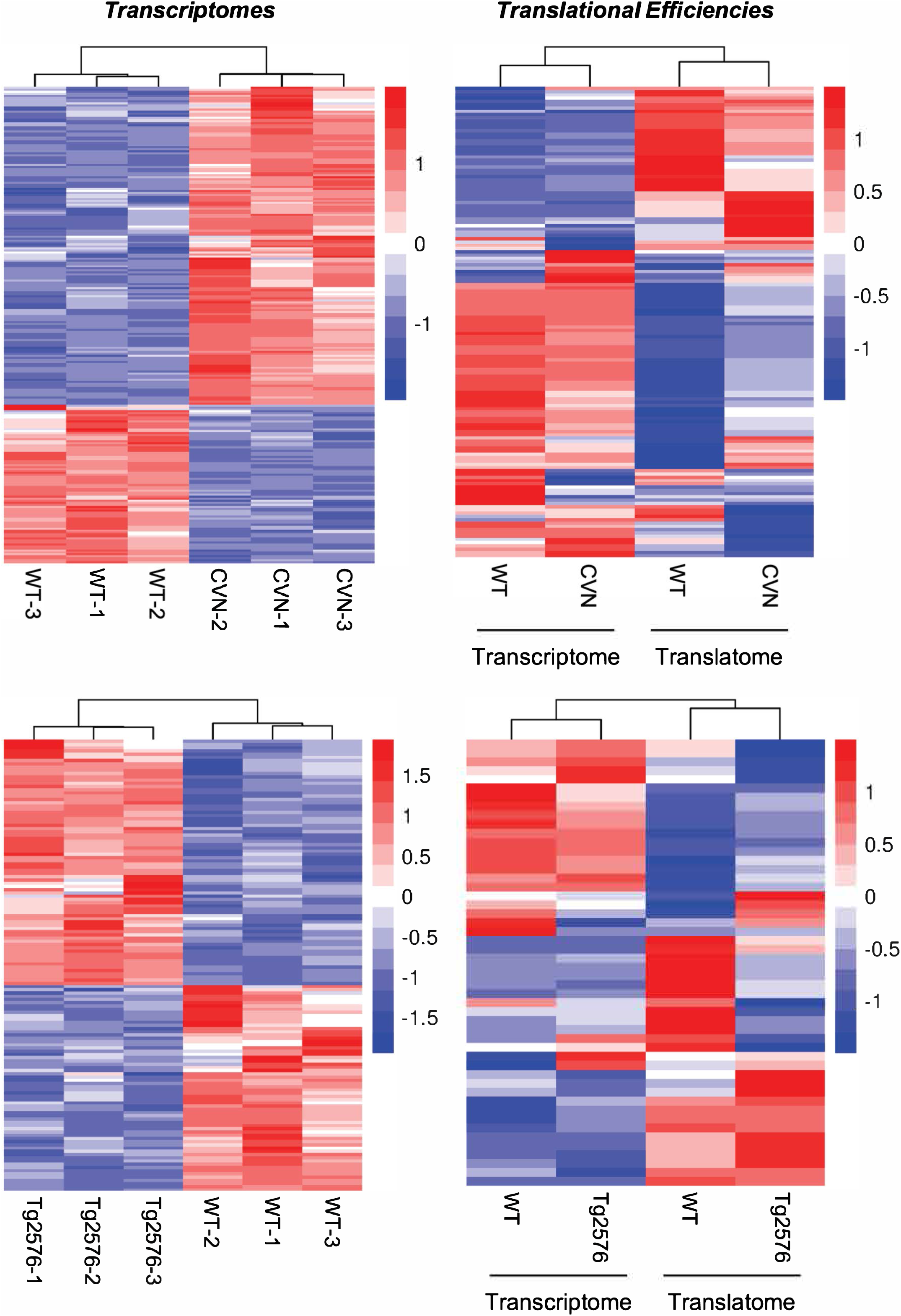
Heatmaps of differentially expressed genes (FC >1.5 and p-adjusted value <0.05).

### Regulation of distinct biological pathways in cortices of CVN versus Tg2576 mice

We next used Ingenuity Pathway Analysis (IPA) [36] to identify biological functions associated with genes that were up-regulated or down-regulated at FC >1 in CVN and Tg2576 mice compared to their respective WT counterparts. CVN mice showed a clear functional response at the transcriptome, translatome and translational efficiency levels (Fig. 5, S10 and Table S4). Down-regulated biological functions in CVN mice were dominated by processes indicative of neuronal decline, such as neurodegeneration, demyelination, and decreased axon growth, LTP and glutamate release. In contrast, indicators of myelination, glial cell abundance and neurotransmission were identified by IPA as major up-regulated processes in CVN mice. IPA also detected an up-regulated neuroprotective response for microglia, which may reflect the early accumulation of Aβ. However, IPA also detected activation of neurodegeneration-related processes like reduced axon growth and increased neuron death, especially at the transcriptional level (Table S4). Moreover, at the translational efficiency level, IPA reported a marginal (low z-score) decrease in LTP related genes (Fig. 5).

**Fig. 5.**
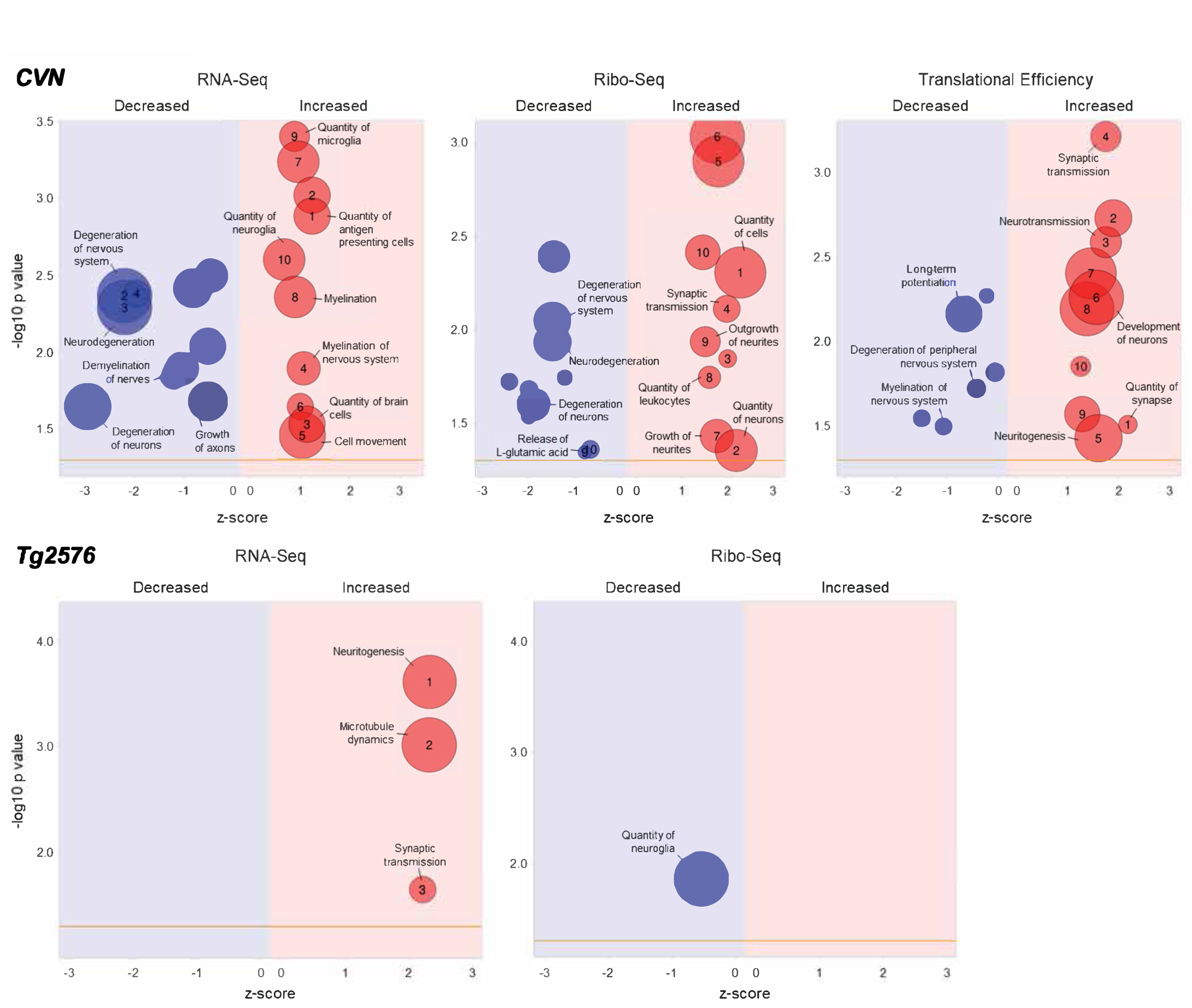
Functional enrichment analysis of differentially expressed genes in CVN and Tg2576 mice. The top 10 (z-score) decreased and increased functional categories are shown for RNA-Seq, Ribo-Seq and Translational Efficiency. Analysis was performed using Ingenuity Pathway Analysis and graphical representation was obtained from *GOplot* R package. Table S4 contains the complete set of decreased and increased pathways at each level.

In contrast, only a few overrepresented functional categories in Tg2576 mice were identified with IPA (Fig. 5 and Table S4). At the transcriptome level no categories were found to decrease, and only three categories increased: neuritogenesis, microtubule dynamics and synaptic transmission. At the translatome level only the category of quantity of microglia was decreased, but with a low z-score, and no categories increased. No functional categories in translational efficiency were detected.

Using IPA again, we then performed an upstream analysis to infer modulated genes by the observed regulation of their targets. For CVN mice, most of the upstream genes were closely related to AD pathology and implied protective reactions to avoid Aβ accumulation and counter neurodegeneration (Fig. 6). For example, upstream genes that were suppressed included: *PSEN1*, *PSEN2*, *APOE*, *B4GALNT1*, *ST8SIA1* and *CTCF* at the transcriptome level; *SSB* at the translatome level; and *ADORA2A* at the translational efficiency level. Decreased expression levels of *PSEN1, PSEN2* and *APOE* could signal a defense response against Aβ production, as might reduced expression of *B4GALNT1* and *ST8SIA1*, ganglioside synthases that increase APP cleavage and affect memory [41, 42]. *CTCF* encodes a transcription factor that can act as an activator or repressor, and also serves as an insulator protein that defines chromatin domains and can up-regulate APP expression [43]. *CTCF* knockout in mouse hippocampus increases cytokine expression and activates microglia [44]. *ADORA2A* is an adenosine receptor, and its pharmacological inhibition or downregulation restores LTP and reverses memory deficits [45, 46]. On the other hand, upstream genes indicated as activated seem to represent both protective and degenerative responses (Fig. 6). *TCF7L2* and *CSF1*, which we found to be activated at the transcriptome level, probably have a protective function. *TCF7L2* may be involved in improving neurogenesis and compensate neuron loss [47] and CSF1 has been linked to microglial activation, prevention of cognitive loss and reduction of Aβ accumulation [48, 49]. *PTF1A* and *IFNG*, identified as upstream activated genes at the translational level, may also be protective. PTF1A can induce neuronal stem cell generation and improve cognitive dysfunction [50]. *IFNG*, which encodes interferon gamma (IFN-γ), can activate microglia to suppress Aβ deposition and induce neurogenesis [51–53]. In contrast, other genes found to be activated, such as *TNF*, may promote degenerative processes. *TNF* encodes tumor necrosis factor alpha (TNF- α), a proinflammatory cytokine that exacerbates both Aβ and tau pathologies *in vivo* [54]. However, the roles of cytokines like IFN-γ and TNF-α are controversial in AD [55].

**Fig. 6.**
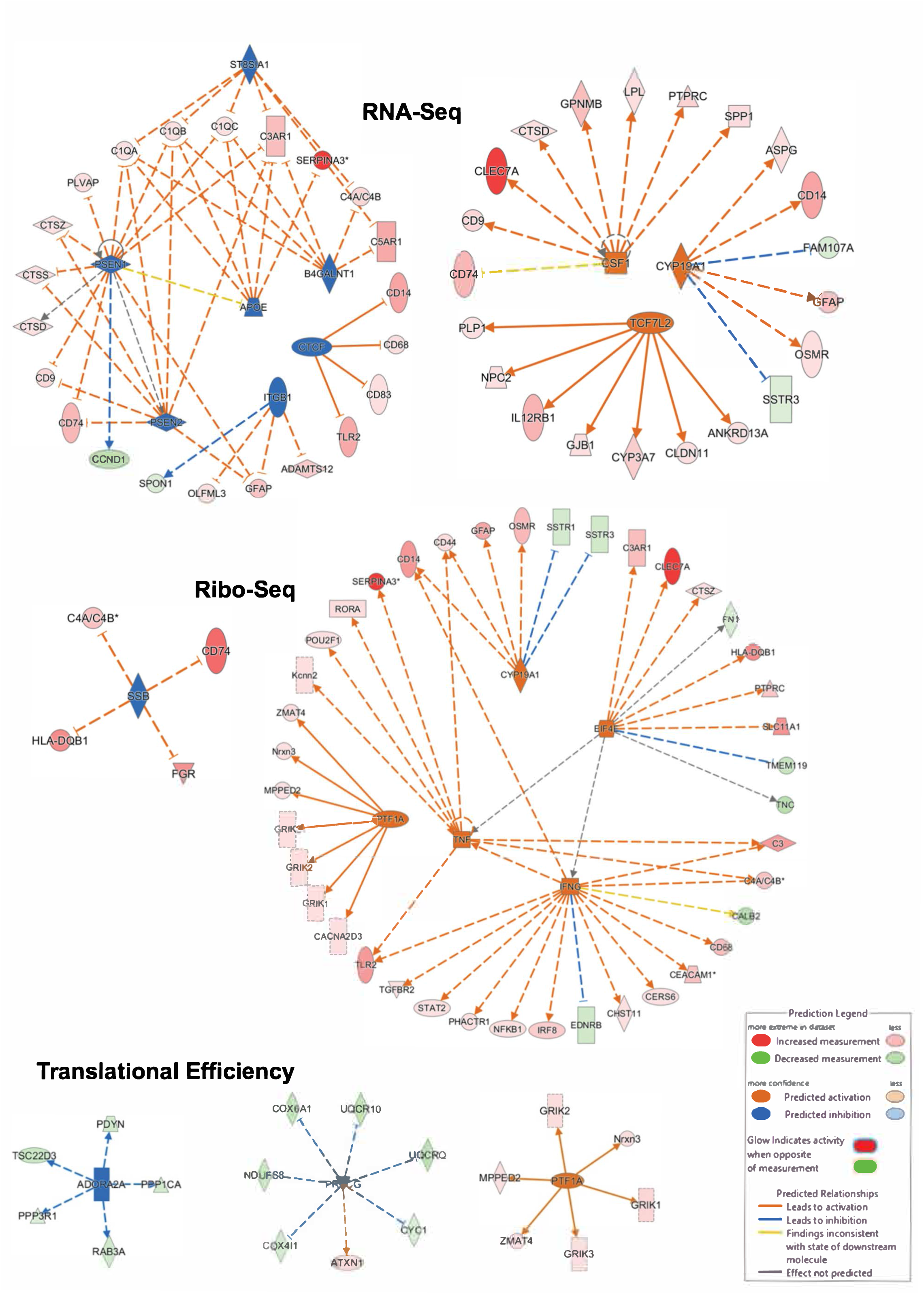
Upstream regulation in CVN mice predicted from differentially expressed gene lists by Ingenuity Pathway Analysis.

The upstream activated genes identified from the translational efficiency DEGs, *PTF1A* (see above) and *PRKCG*, are both probably protective. Protein kinase C γ, which is encoded by *PRKCG*, can stimulate APP processing by α-secretase to produce soluble fragments of APP and reduce Aβ accumulation [56]. In contrast, we did not detect any prominent upstream gene regulation in the Tg2576 mice as might be expected from the uninformative functional enrichment analysis. Collectively, our functional analyses reinforce the conclusion that the CVN and Tg2576 models exhibit differential cortical brain gene regulation at six months of age.

### Regulated genes in CVN mice reveal a signature of microglia responding to Aβ

To extend the functional analysis further, we used *IdMiner,* a tool that we developed in-house (manuscript in preparation; https://github.com/sradiouy/IdMiner) to enhance interpretation of DEGs. This software is a text-mining tool that captures previously reported associations between gene lists and user-defined terms using the PubMed database. We found that among up-regulated genes in CVN mice at both the transcriptome and translatome level there was a clear signature of disease-associated microglia [20]. This signature included genes such as *TREM2, TYROBP, TLR2, CD68, GPR84, GPNMB, ITGAX, ITGB2, LPL, CLEC7A, CST7* and *CCL6* [20,21,57–60]. All these genes encode transmembrane proteins that are highly expressed in a microglial subpopulation that specifically responds to neurodegeneration. Overexpression of these markers orchestrates clearance of Aβ by a subpopulation of disease-associated microglia [20,57,61]. This signature is part of the immune system response in AD, a statistically overrepresented function among up-regulated gene networks at both the transcriptome and translatome levels (Fig. S11 and S12).

Other genes up-regulated at both the transcriptome and translatome levels included those expressed primarily either in non-neuronal or neuronal cells, but not in both. For example, *GFAP* and *SERPIN3* are overexpressed in astrocytes in AD [62–65]. Solute carrier family members and hippocalcin-like proteins were also up-regulated, in accordance with their possible neuroprotective functions in AD [66, 67]. Additionally, we found increased expression of *MID1*, which encodes a member of the MID1 protein complex that binds to and accelerates *APP* mRNA translation through an mTOR-dependent pathway [68]. The tyrosine kinase gene, *FGR,* was also up-regulated at both transcriptional and translational levels. Fgr is able to bind tau, and although functional consequences of this interaction are not well understood [69], the Fgr paralog, Fyn, binds and phosphorylates tau [70] causing neurotoxicity [71, 72].

An analysis of the genes up-regulated specifically at the translatome level revealed other examples of AD-associated genes. Examples include the histocompatibility 2 class II antigen A gene, *H2-AB1*, a marker for disease-associated microglia [57, 58], and complement components 4a and 4b, which were up-regulated as previously described in response to Aβ plaque increase [65]. The *CSF3R* gene was also up-regulated at the translatome level, as expected, since it is highly expressed in early stages of AD [73]. Deficiency in G-CSF, a ligand for the receptor encoded by *CSF3R*, has a deep impact in hippocampal structure and function leading to disruption in memory formation and impaired behavioral performance [74]. Likewise, injected G-CSF acts as a neurotrophic factor and induces astrocytic and microglial activation [75, 76], so elevated *CSF3R* expression may signify a response mechanism to mitigate neurodegeneration.

When studying the list of up-regulated genes particularly at translational efficiency level, we detected several notable genes: *SERPINA3G, FRMD4A, RRAS2, AV2, ADAM12, PPARG, MID1, LDLRAD3* and *CD44*. Some of these genes are associated with protective or beneficial functions. For example, *FRMD4A*, *PPARG* and *RRAS2* have been shown to reduce high levels of Aβ production [77–80], while *ADAM12*, *MID1* and *LDLRAD3* influence Aβ production or neurotoxicity [68,81,82].

Regarding down-regulated genes, we also found examples related to Aβ accumulation. *CD59*A (complement defense 59A) was down-regulated at the transcriptome level, as might be expected in response to Aβ, and could contribute to neuronal vulnerability and loss [83, 84]. Calbindin 2 (calretinin; CALB2) positive interneurons are specifically decreased as early targets of Aβ accumulation in the hippocampus [85] and in the current study its expression was down- regulated at the transcriptome and translatome levels. *PIN4* was also down-regulated at both levels in accordance with the decrease observed for Pin1, a paralog gene, which plays a role in the accumulation of Aβ [86–88] and tau [89, 90]. Other down-regulated genes that may be germane to neurodegeneration include *RPLP0* and *IGF2* [7, 91], while genes like *SLC17A8* and *TNC* (tenascin-C) may have been down-regulated to ameliorate pathology [92].

### Tg2576 mice differentially express genes associated with APP metabolism

The number of DEGs in Tg2576 mice was much less than in CVN mice. The Toll like receptor 6 gene (*TLR6*) was up-regulated at both the transcriptome and translatome levels. Tlr6 can dimerize with Tlr4 and interact with CD36 to bind Aβ and activate microglia, and thereby promote neurodegeneration [93]. Calpain 11 was also up-regulated at the transcriptome level, as expected since calpains are hyperactivated in AD [94, 95]. CDK5RAP1, an inhibitor of Cdk5, was also up-regulated and could have a protective effect since Cdk5 activation causes hyperphosphorylation of APP and tau leading to plaques and tangles [96–98]. Also, Cdk5 has been described as a mediator of Aβ-induced neuronal cell cycle reentry that leads to neurodegeneration [99].

At the translatome level, evidence of other interesting up-regulated genes was found. Apolipoprotein D, which has a neuroprotective effect [100] and is induced in hippocampal cells in response to Aβ [101], was translationally increased. Another gene with a neuroprotective function up-regulated at the translatome level encodes Metallothionein 3, which is expressed by astrocytes and is thought to facilitate Aβ uptake [102], although this role remains controversial with opposite findings *in vitro* and *in vivo* and with different mouse models. For example, a study using the Tg2576 mice model shows that Metallothionein 3 could have opposite effects depending on gender, brain region and age [103]. Genes up-regulated exclusively at the level of translational efficiency were related to Aβ production and disease. Examples include *SERF2*, a positive regulator of amyloid protein aggregation [104, 105], and *HRAS2*, an apparent AD biomarker that is stimulated by Aβ and produces a reduction in LTP [78].

Some transcriptionally down-regulated genes confirmed what was expected from previous reports. For example, *KLK6* encodes a peptidase that cleaves APP and is down-regulated in the cortex of human postmortem samples of AD patients compared to controls [106, 107]. The somatostatin receptor 5 was down-regulated at both the transcriptome and translatome level, consistent with the decreased levels reported for these receptors in the cortex of AD patients [108, 109]. The *FABP5* gene was found to be down-regulated at the translational efficiency level as expected since a Fabp5 deficiency is associated with increased vulnerability to cognitive deficits in mice with APP pathology [110, 111]. Thus, Tg2576 mice showed differentially expressed genes associated with APP metabolism at both the transcriptome and translatome level.

### Regulated functions common to both CVN and Tg2576 mice are involved in APP metabolism

We next compared the DEGs detected for CVN and Tg2576 mice using a FC cutoff of 1 to explore commonalities shared by both mouse models (Table 2). Common genes were rare, representing <10% of the genes in our DEG list. When comparing up-regulated genes at the transcriptome level between CVN (282 genes) and Tg2576 (195 genes) mice, we found 25 shared genes. Of those, 19 are classified as predicted genes. Exhaustive analysis revealed that those genes mainly encode Zinc finger and α-takusan-like proteins, but they also include a long non-coding RNA (*Gm26650*). α-takusan proteins represent a large family that regulates synaptic activity and reportedly can mitigate Aβ-induced synaptic loss [112, 113]. Among the common and annotated genes, we found *LRRC37A*, a leucine-rich repeat containing gene that encodes a plasma membrane protein involved in intracellular vesicle trafficking [114] and *OSMR* (Oncostatin M Receptor). *LRRC37A* was previously associated with AD because of its many SNPs detected in genome-wide association studies of *APOE4* carriers and its genomic location adjacent to the tau gene, *MAPT* [115]. On the other hand, it has been reported that Oncostatin M is neuroprotective against Aβ toxicity [116].

When exploring the translationally up-regulated genes shared by both transgenic models, several interesting examples were noticed. Due to the small number of such genes in Tg2576 mice (66 genes, Table 2), we only found 6 genes in common. However, among them we found *LRRC37A* and a predicted gene (*GM3173*) annotated as an α-takusan-like protein (see above). Other interesting genes found were *GFAP,* and Hemoglobin alpha 1 and 2. Glial fibrillary acidic protein (GFAP) is an intermediate filament protein highly expressed in the reactive astrocytes that surround Aβ plaques [62, 117]. Hemoglobin alpha 1 and 2 (encoded by *HBA-A1* and *HBA-A2*), is expressed by neurons [118], and can bind Aβ and co-localize with plaques [119, 120].

We found 13 down-regulated genes common to CVN and Tg2576 mice, 8 of which were down-regulated at the transcriptome level and 5 of which were down-regulated translationally. Here we highlight *TMEM59L*, which was down-regulated transcriptionally and encodes transmembrane protein 59-like, an important paralog of *TMEM59*. TMEM59 has been identified as a novel modulator of APP shedding, controlling APP post-translational modification, trafficking and cleavage [121]. More recently, it has been reported that TMEM59 interacts with TREM2 (see above and Discussion), and that TMEM59 homeostasis is regulated by TREM2 in order to control microglia activity [122]. Common genes down-regulated at the translatome level include ribosomal proteins and ribosome biogenesis factors (Rpl5, Rpl32 and Wbscr22), the fatty acid binding protein 5 (FABP5) and adaptor related protein complex 4 (AP4S1). It has been reported that the adaptor protein 4 interacts with APP, and that the disruption of this interaction stimulates APP cleavage and Aβ production [123].

Finally, despite the low identity overlap of DEGs in CVN versus Tg2576 mice, we also observed biological functions shared by the two AD mouse model strains. For example, several genes affected in each strain are associated with APP and Aβ metabolism, either with neuroprotective or neurotoxic functions (Fig. 7).

**Fig. 7.**
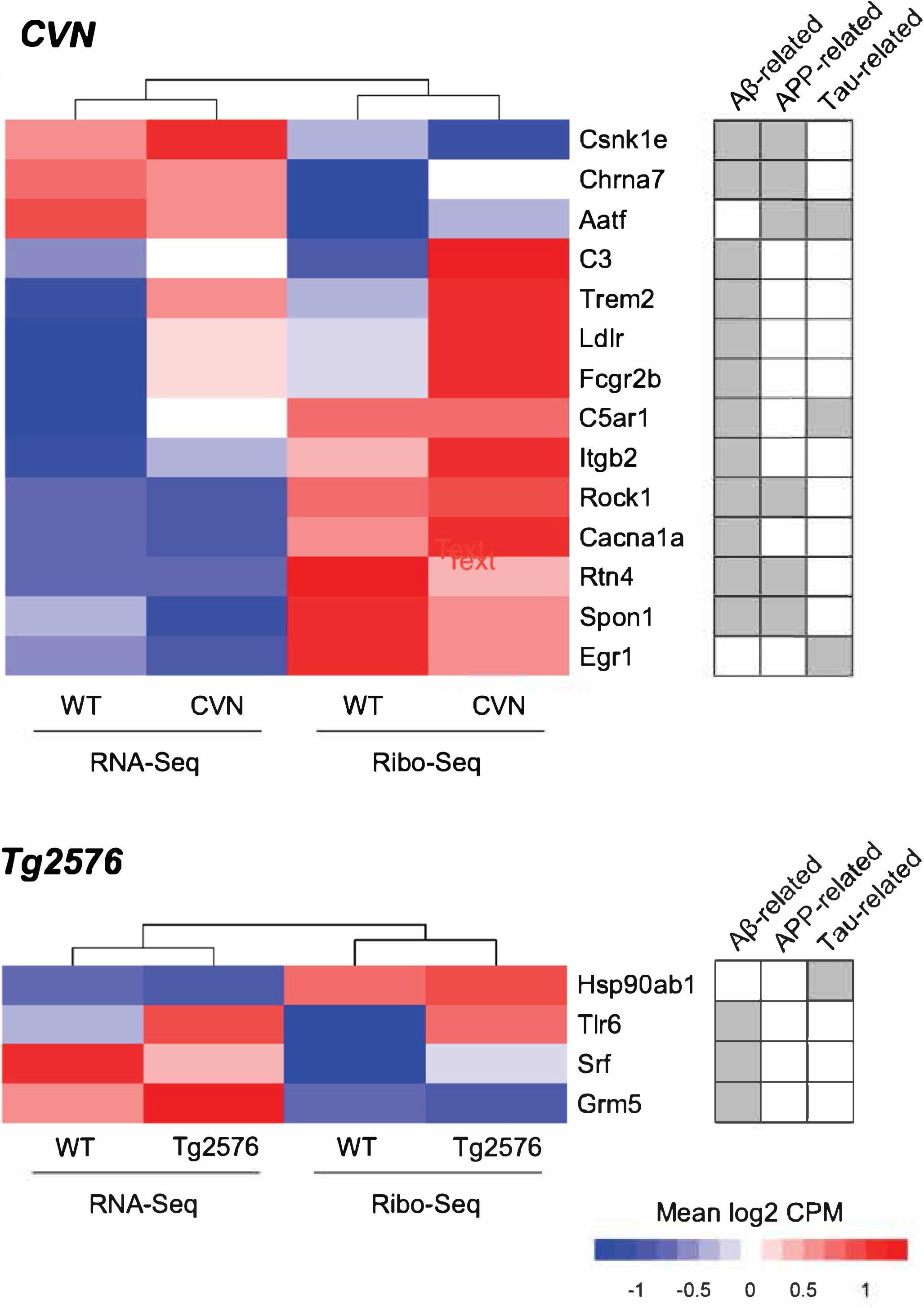
Expression regulation of genes related to Aβ, APP or tau metabolism. Several gene ontology categories obtained from AmiGO2, were merged to consider all genes related to Aβ, APP or tau metabolism. Genes were selected by p-adjusted value <0.05 and the expression levels of those significant genes are shown on the heatmaps.

## DISCUSSION

We used RNA-Seq and Ribo-Seq to explore transcriptional and translational gene expression regulation, respectively, in the brain of two transgenic mouse models of AD. Studies of translation regulation using Ribo-Seq in mouse brain tissue have been sparse (see examples in [124–128]), and, to our knowledge, our work represents the first approach in AD models. Both transgenic mice used in this study are amyloid models since each strain has a transgenic insertion of the APP gene with specific mutations that increase levels of Aβ and amyloid plaques (Table 1). Animals were euthanized at 6 months of age, when CVN mice are asymptomatic both behaviorally and histopathologically (Fig. 1 and <u>CVN - Alzforum</u>) but Tg2576 mice exhibit synaptic loss in the CA1 region of the hippocampus, reduced long term potentiation in the dentate gyrus, and a variety of cognitive impairments (Fig. 1 and <u>Tg2576 - Alzforum</u>).

The isolated total RNA and ribosome footprints obtained from the brain cortex (Fig. S1) yielded more than 20 and 120 million paired-end and single-end reads, respectively (Table S1). We specifically used the brain cortex, a region highly degenerated and pathologically compromised in both humans with AD and murine models of the disease. As anticipated, around 80% of the ribosome footprints samples were rRNA fragments derived from the RNAse digestion. We therefore resorted to deep sequencing to obtain workable amounts of non-rRNA reads. By doing so, we obtained more than 10 million ribosome footprints mapping over mRNAs in each sample (Table S1) and we defined expression levels for more than 14 thousand genes in each transgenic strain. As expected, almost all genes that we detected were found at both the transcriptome and translatome levels (Fig. S2). By studying mapping features and read periodicity, we separated transcriptome-derived from translatome-derived reads (Fig. S4 and S5). For example, translatome-derived reads show the classical three nucleotide mapping pattern periodicity not detected in transcriptome-derived reads, providing evidence of solid transcriptome and translatome datasets.

Differential gene expression analysis was performed over transcriptome and translatome samples separately, comparing transgenic mice versus WT, to explore deregulated genes in each model. However, considering that differences at the translatome level could be explained by regulation at the transcriptome level, we estimated translational efficiency as the ratio between translatome and transcriptome levels to specifically identify translational regulation events [23] (Fig. 2 and 3). Despite using different animals as replicates, which could generate considerable biological variation, samples clustered as expected as evidenced by principal component analysis (Fig. S6). Also, we were able to define lists of genes with a significant statistical difference between genotypes at p-adjusted values <0.05 (Table 2). Expression levels of DEGs also separates samples as expected and shows the different levels of regulation observed, especially when analyzing translational efficiency regulation (Fig. 4). As a control comparison, we also explored differential gene expression between WT animals, and there is only a limited overlap with previously defined DEGs and the enriched biological functions observed were either not related to the results previously discussed or absent (Fig. S13 and S14).

The functional analysis of regulated genes involves different approaches and techniques. We used standard gene ontology and functional enrichment tools, namely *IPA* and *STRING*, which revealed a clear signature of decreased neurodegenerative-related process and increased neuroprotective-related functions, particularly for the CVN strain (Fig. 5 and 6). Secondly, we complemented the analysis with an in-house developed tool (https://github.com/sradiouy/IdMiner) to deeply explore DEGs lists in order to find relevant associations between genes and biological functions related to pathology. By using this approach, we detected several genes associated either with the neurodegenerative and neurotoxicity phenotype or with protective and beneficial functions (see below).

In CVN mice we identified a clear gene signature common in disease-associated microglia (DAM). These genes mainly encode transmembrane proteins that were up-regulated at both transcriptome and translatome levels, and mediate immune system processes (Fig. S11 and S12). Earlier studies of other AD mouse models at the same or older ages suggest that the main functions of this DAM population are to locate and clear Aβ plaques [20,21,57,58]. It is noteworthy that a DAM population is present in CVN mice at a stage when they do not show any phenotypic signs of disease. Among the genes present in this signature are *TREM2* and *TYROBP,* both of which were up-regulated (transcriptome level: FC = 2.12 and 1.66, p-adjusted value <1.18E-15 and <6.22E-4, respectively; translatome level: FC = 1.90 and 1.72, p-adjusted value <4.25E-8 and <1.39E-4, respectively). The TREM2/TYROBP complex expressed in microglia is necessary to prevent Aβ accumulation and diffusion [20,129–131]. TREM2/TYROBP-dependent cell activation seems to be beneficial [132], while *TREM2* deficiency impairs cellular metabolism and promotes increased autophagy in microglia in an AD mouse model [133]. Concordantly, in CVN mice we found several members of the TYROBP AD-related pathway [59, 134] regulated at the transcriptome and/or translatome level (Fig. S15). Regarding *TREM2*, transgenic overexpression in an AD mouse model of the human version of this gene modified the morphological and functional responses of microglia, which resulted in amelioration of the pathology and memory deficits [135]. Interestingly, it has been reported that TREM2 interacts with TREM59 and modulates microglia activation [122]. Here we found *TREM59L* as a down-regulated gene at the transcriptional level in both transgenic mice models (CVN: FC = -1.89 and p-adjusted value <3.56E-28; Tg2576: FC = -1.29 and p-adjusted value <0.01). Regarding functional interpretation of DEGs and considering the upregulated genes at the translational level, two different sets of genes were observed. One set associated with protective functions, such as reducing Aβ production, and others related to the high levels of Aβ and neurotoxicity. A similar interpretation was made from the lists of down-regulated genes.

Although Tg2576 mice experience cognitive deficits at 6 months, amyloid plaques typically do not appear in this strain until 11-13 months (Fig. 1 and <u>Tg2576 - Alzforum</u>). Nevertheless, we found a group of genes that were regulated at the transcriptome and/or translatome levels in the 6 month old Tg2576 mice, and have been reported to be altered in response to the presence of Aβ. In light of the dearth of plaques, we suspect that expression of these genes was affected by accumulation of Aβ oligomers in cortical brain tissue. Also, as observed for CVN mice, we found genes associated with neuroprotective functions and others with a neurodegenerative phenotype related to APP metabolism (Figure 7).

We also explored DEGs that were common to both AD strains at the transcriptome or translatome levels. Despite the facts that CVN and Tg2576 mice are derived from different WT parental strains (Table 1), and exhibit different patterns of phenotypic alterations (Fig. 1A) and distinct regulation levels (Table 2), it is noteworthy that they share genes that are regulated transcriptionally and translationally. These DEGs, such as α-takusan proteins, *LRRC37A*, *OSMR*, *GFAP*, hemoglobin alpha 1 and 2, *TMEM59L* and *FABP5*, are seen at early stages of disease and modulate functions that could be associated with AD pathology. Despite the lack of plaques in 6 month old CVN mice, most of the DEGs common to both strains are implicated in APP metabolism in a neuroprotective fashion that responds to Aβ accumulation.

It is important to note that we analyzed gene expression of a complex brain tissue that contains a heterogeneous population of cells. In addition to specific regulation events occurring in particular cell types that could be underestimated here, the high abundance of neurons adds an extra layer of complexity, because of the large dimensions and high polarity these cells can achieve. Considering the relevant contribution of local regulation events in neurons [136–139] and in particular previous evidence in the context of AD [11], we cannot exclude the possibility that local protein synthesis events may regulate neuron response to local stimulus in the brain cortices. To specifically disentangle these regulatory events, one would need to conduct new experiments using tools designed to examine functional genomics in specific neuronal compartments [17].

Overall, this study documents the contribution of the different layers of gene expression regulation involved with AD pathology, especially at the level of translation. The two amyloid-producing animal models used here show relevant gene expression regulation at the transcriptome and translatome levels. Although the number of differentially expressed genes was markedly different, both models reveal several regulated genes associated with APP and Aβ metabolism. However, we noticed that each transgenic mouse was able to modulate different groups of genes that represent specific biological functions. For example, although the 6 month old CVN mice that we used were asymptomatic for AD-like traits, their gene expression profiles were skewed towards neuroprotection. Degeneration processes were decreased, protective functions were increased, and a specific microglia subpopulation was induced as a presumptive response to Aβ accumulation prior to plaque formation. On the other hand, 6 month old Tg2576 mice displayed signs of simultaneous neuroprotection and neurodegeneration. Perhaps this reflects a more rapidly progressing AD-like phenotype manifest as cognitive impairment at that age for Tg2576 mice (<u>Tg2576 - Alzforum</u>), as compared to ∼13 months for CVN mice (<u>CVN - Alzforum).</u>

Recent advances in genomic views of translation have been essential to widen our current concepts about the dynamic properties of protein translation as a process. More specifically, such advances increase the relevance that regulation of translation has on particular sets of mRNAs, and by extension on proteostasis. Besides pointing to novel transcriptional regulation events in AD disease models, here we provide evidence that translation is important in regulating key AD related genes at early asymptomatic stages. Several of the dysregulated pathways observed here will need further study to elucidate regulation mechanisms, but their identification implies that both translation and transcription are dysregulated during both pre-symptomatic and symptomatic phases of AD.

## ACKNOWLEDGMENTS

We thank members of Sotelo-Silveira lab for the results discussion and intellectual input. We also thank members of Bloom lab, especially Drs. Andrés Norambuena and Antonia Silva, for their support with wet lab experiments, and Dr. Dora Bigler-Wang and Nutan Shivange for handling mice and brain dissections. We also acknowledge the following sources of financial support: PhD fellowships from Agencia Nacional de Investigación e Innovación (ANII) to G.E. (POS_NAC_2016_1_129959), Programa de Desarrollo de las Ciencias Básicas (PEDECIBA) to G.E. and J.R.S-S., PROLAB travel grant (PABMB/ASBMB/IUBMB) to G.E.; the Owen’s Family Foundation, the Cure Alzheimer’s Fund and the Rick Sharp Alzheimer’s Foundation to G.S.B., J.S.L. and E.R.S., and NIH grant RF1 AG051085 to G.S.B.

## AUTHOR CONTRIBUTIONS

J.R.S-S. and G.S.B. conceived the project, designed and supervised the research. J.S.L. and E.R.S. contributed to the project design and intellectual content. G.E. performed the experiments. G.E. and E.R.S. analyzed data. G.E., J.R.S-S. and G.S.B. wrote the paper. All authors approved the manuscript.

## SUPPLEMENTARY MATERIAL

**Table S1.**
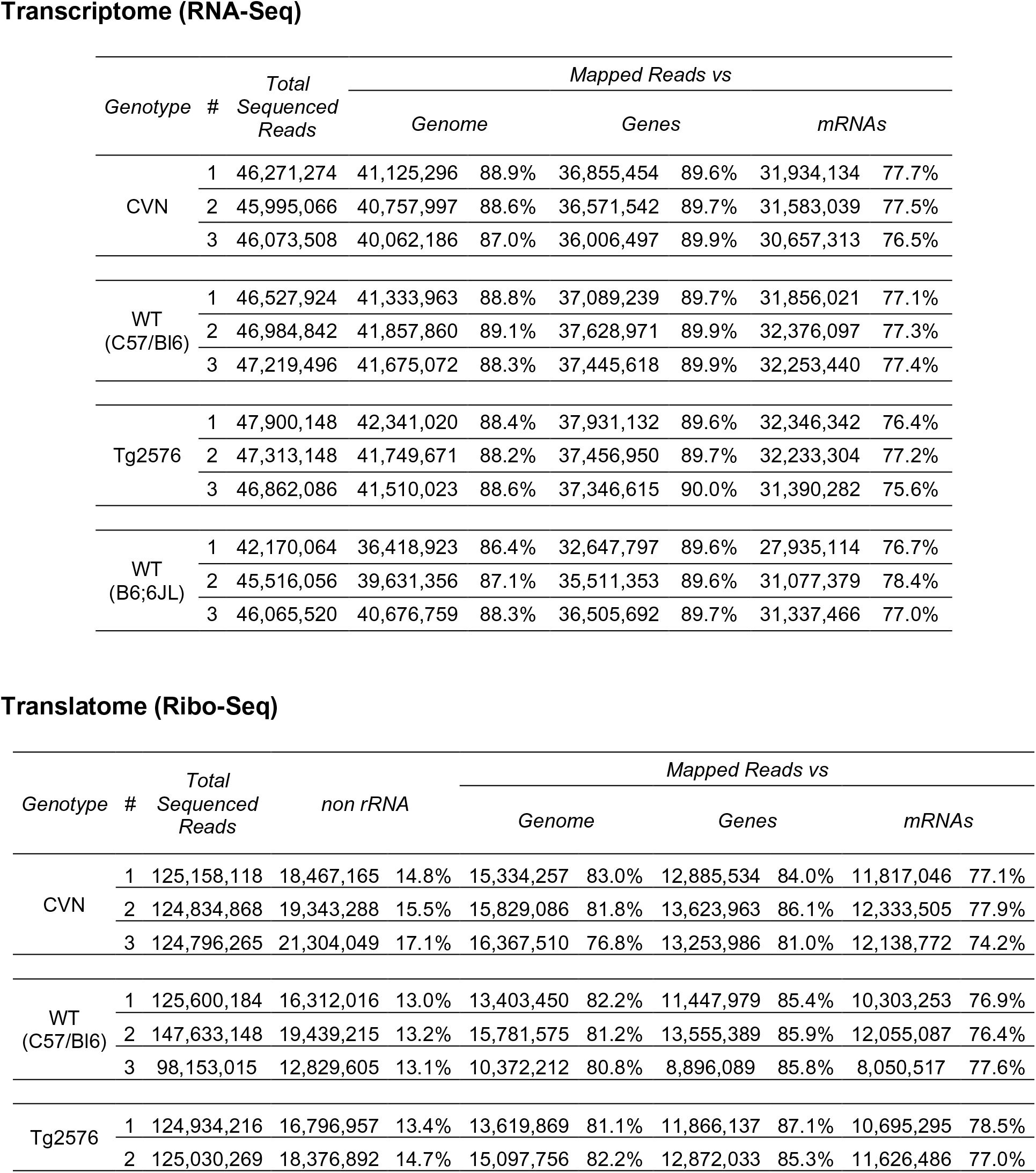
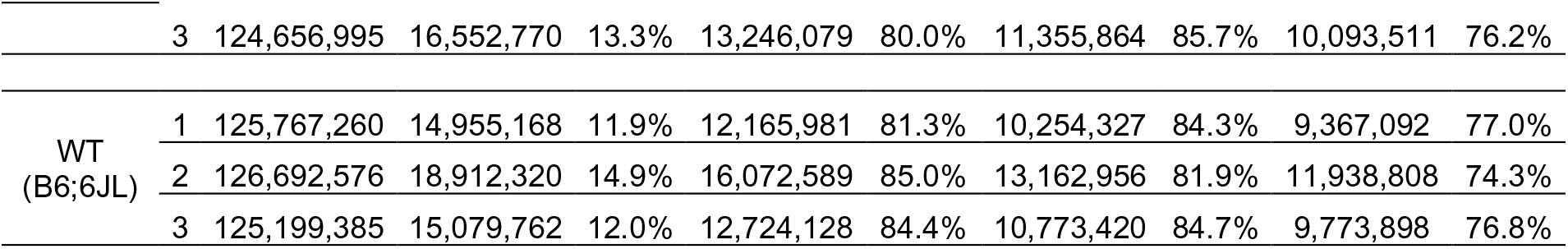
The number of sequenced and mapped reads is shown for each mouse strain for both transcriptome and translatome samples.

Table S2. Differentially expressed genes for CVN mice.

(Excel file attached)

Table S3. Differentially expressed genes for Tg2576 mice.

(Excel file attached)

Table S4. Complete set of decreased and increased pathway for both transgenic mice at the transcriptome, translatome and translational efficiency level determined by *IPA*.

(Excel file attached)

## SUPPLEMENTARY FIGURE LEGENDS

**Fig. S1.**
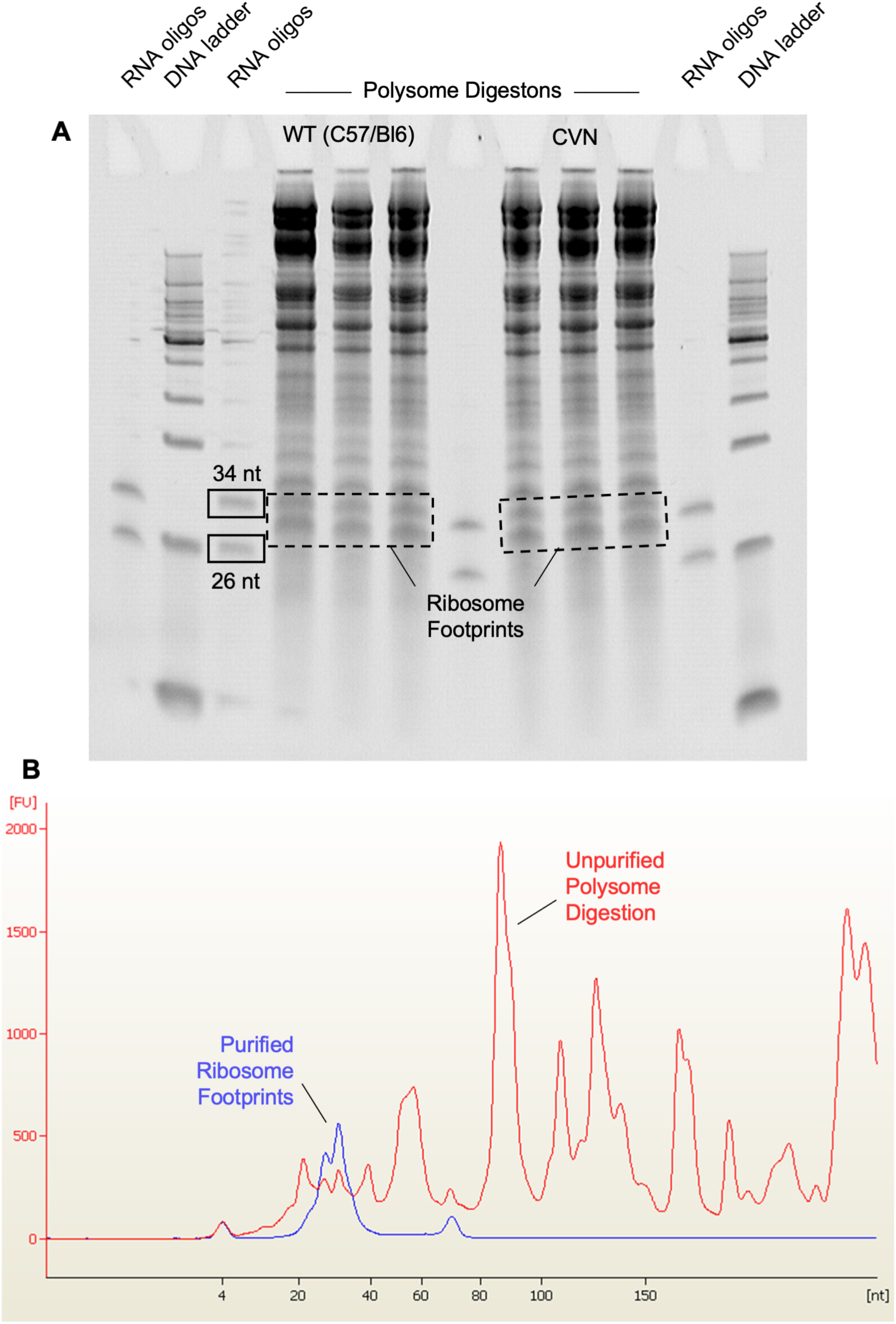
Ribosome footprint purification. A) Polysome fractions digested with Benzonase were separated in a 15% PAGE with 7M Urea. Using RNA oligos as markers, ribosome footprints bands were identified and excised in a dark room under UV light. B) Ribosome footprint quality was evaluated in a 2100 Agilent Bioanalyzer instrument using a Small RNA kit. Representative electropherograms of an unpurified polysome digestion sample and a purified ribosome footprint sample are shown.

**Fig. S2.**
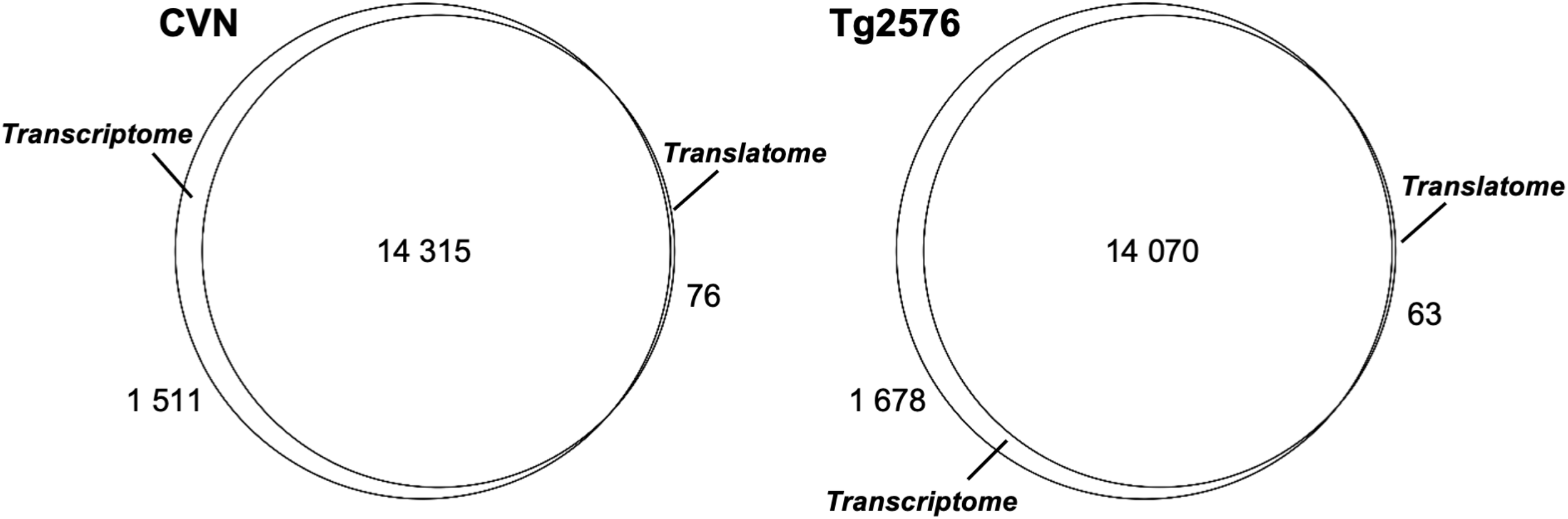
Genes detected by transcriptome (RNA-Seq) and translatome (Ribo-Seq) analysis.

**Fig. S3.**
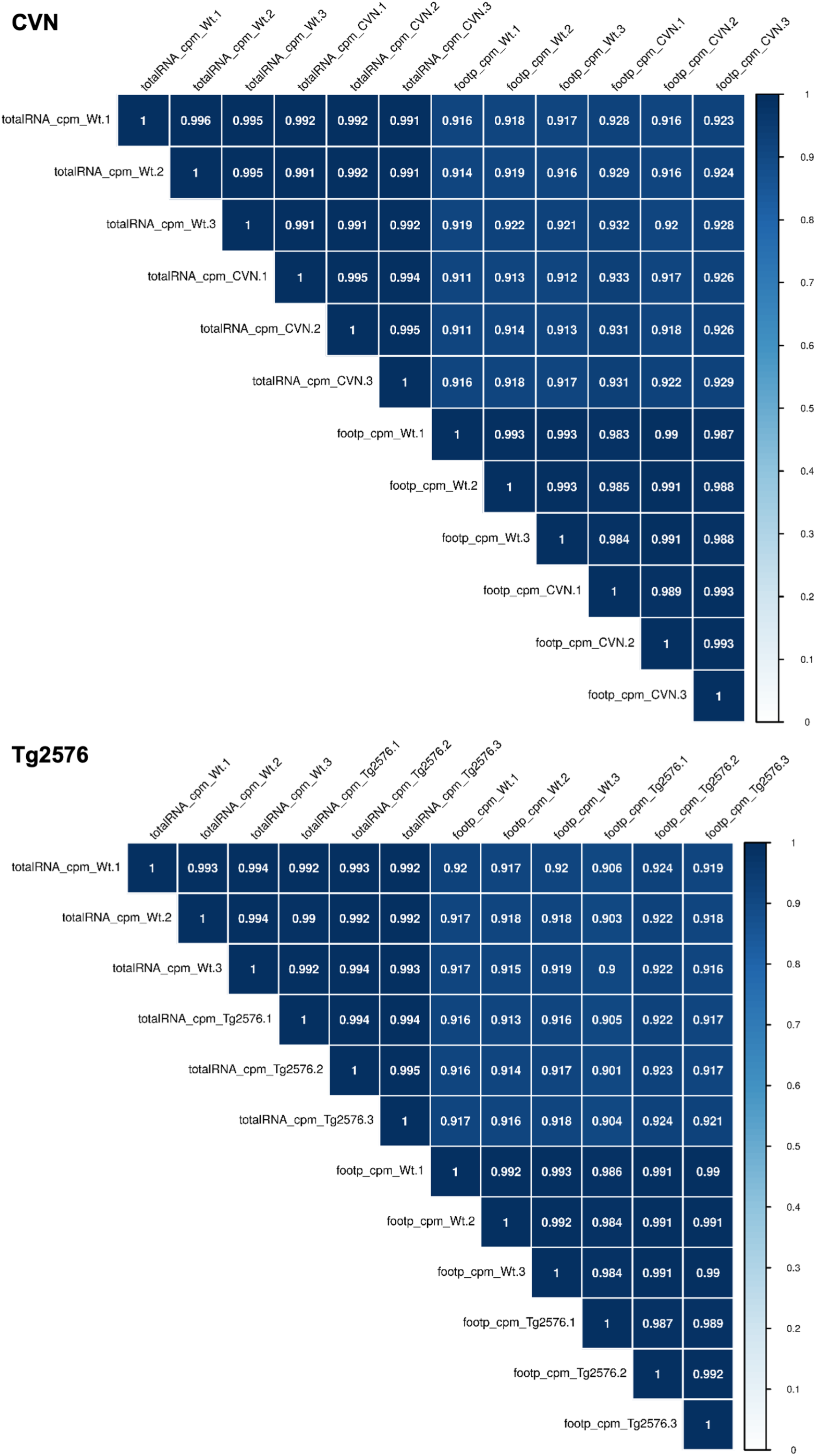
Inter-replicate correlations among transcriptome and translatome samples. Pearson correlation value is shown for each pairwise correlation.

**Fig S4.**
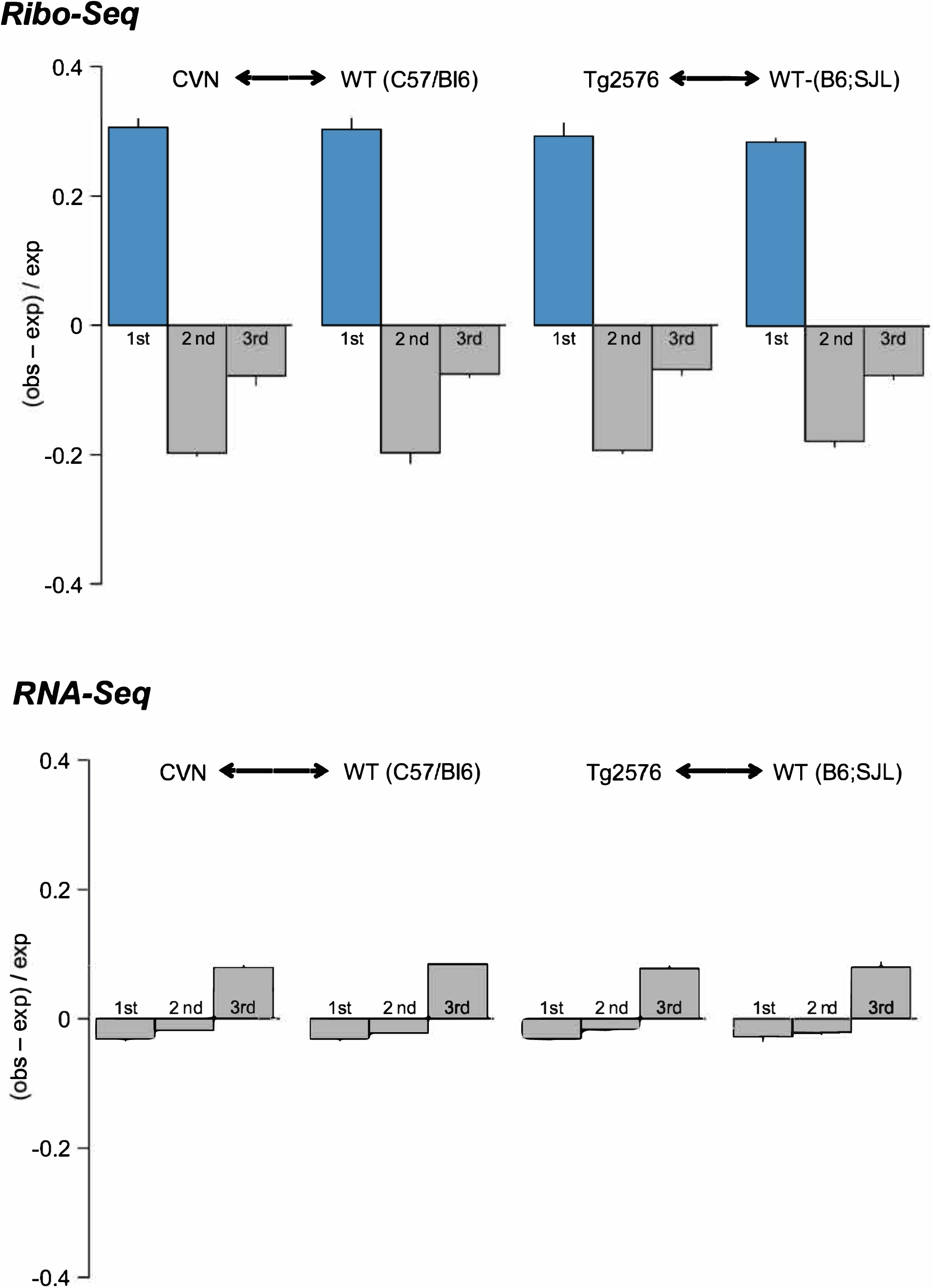
Observed-to-expected ratio of the Ribo-Seq and RNA-Seq derived reads mapping to each of the three nucleotides in codons. Ribo-Seq derived reads 5’ ends are preferentially enriched at the first nucleotide of the codon, while RNA-Seq derived reads 5’ end are distributed more equally among the three nucleotides. This analysis was performed following the criteria described by Ingolia *et al.* Science 2009 and Guo *et al.* Nature 2010.

**Fig. S5.**
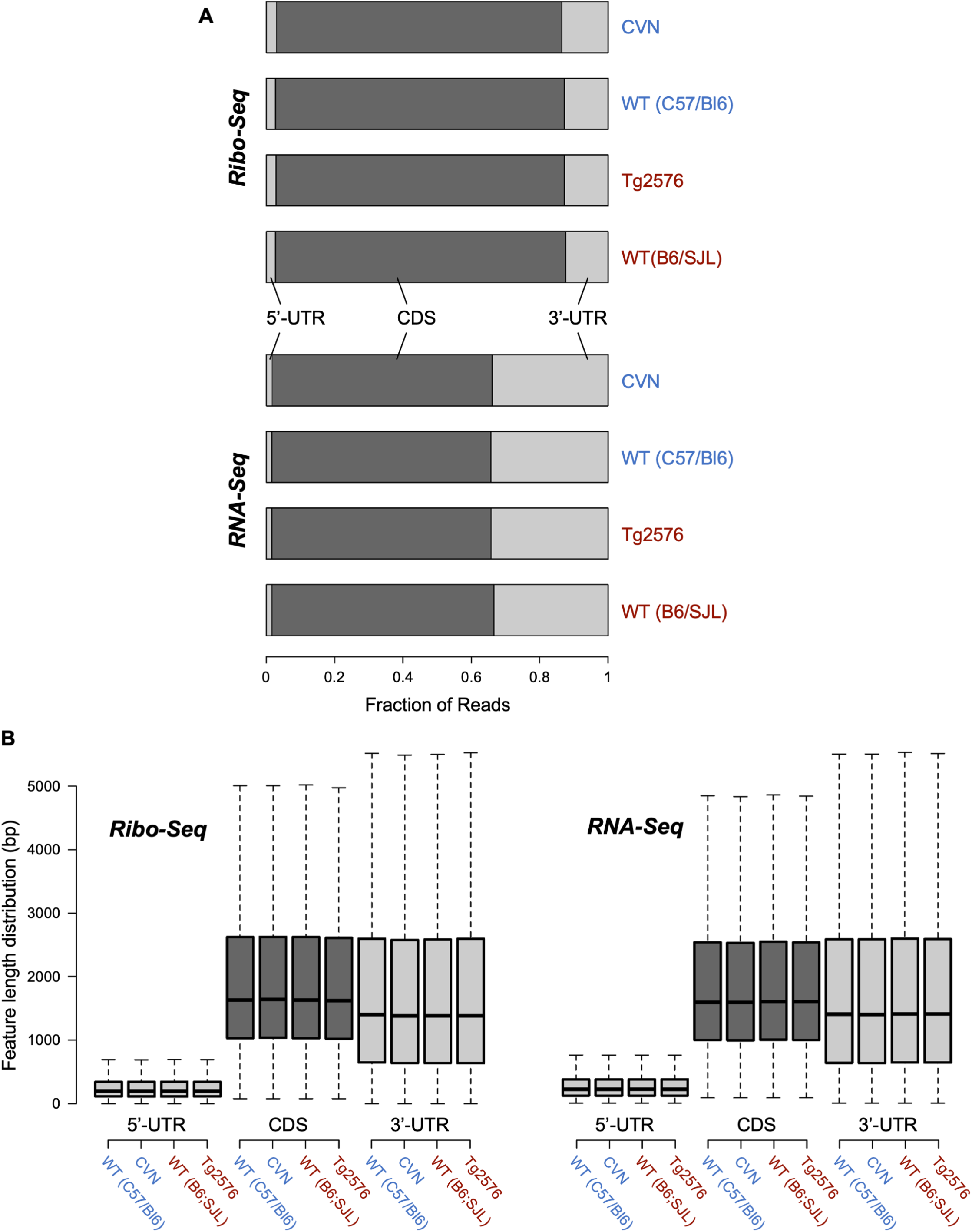
Distribution of mapped reads among coding sequences (CDS), and 5’ and 3’ untranslated regions (5’-UTR and 3’-UTR). A) Fraction of mapped reads among mRNA features is shown for CVN and Tg2576 transgenic mice models and their wild type controls, for the Ribo-Seq and RNA-Seq data sets. Ribo-Seq derived reads map preferentially over CDS and RNA-Seq derived reads are distributed in a more uniform pattern. B) This difference is not due to a bias in feature length.

**Fig. S6.**
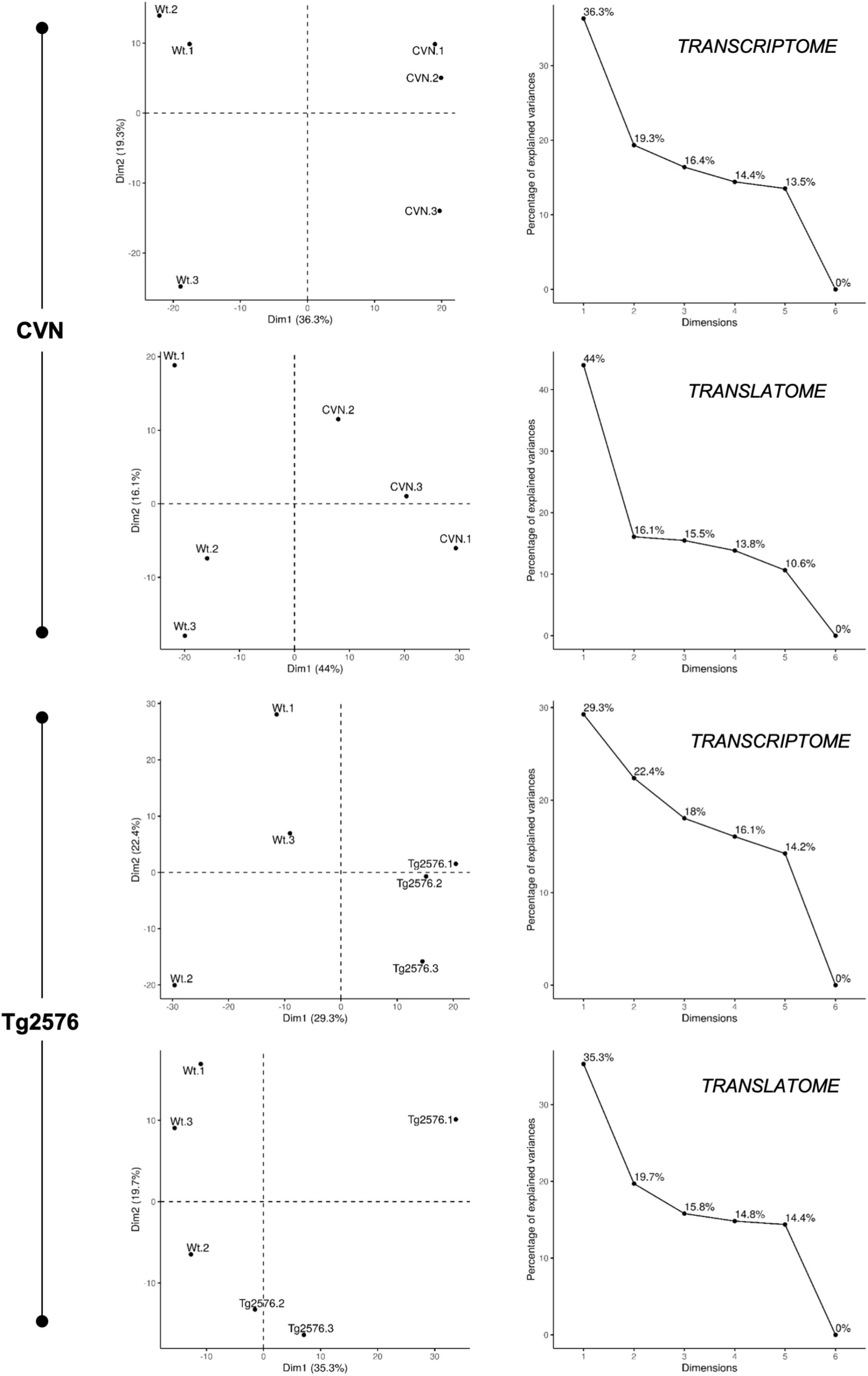
Principal Component Analysis (PCA) for transcriptomes and translatomes. In each case, a 2D plot of principal component 1 and 2 for transcriptomes or translatomes is shown with the scree plots indicating the percentage of explained variation by each dimension of the PCA.

**Fig. S7.**
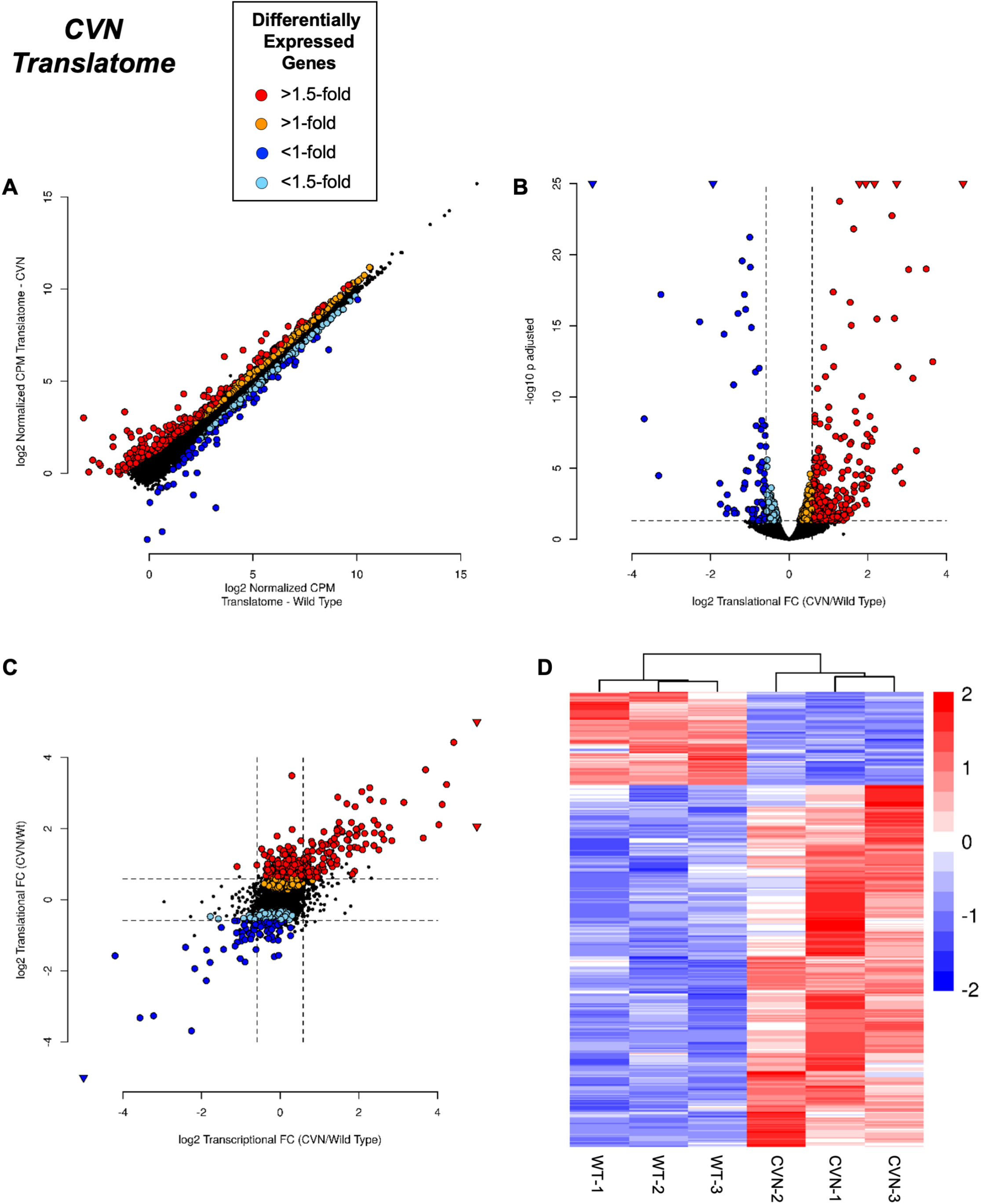
Differential expression analysis performed by *edgeR* for the translatome compartment comparing CVN mice versus their WT parental strain (C57/Bl6). A) Scatter plot comparing normalized CPM expressions at the translational level between genotypes. B) Volcano plot of the relationship between FC and p-adjusted values. C) Scatter plot comparing translational vs transcriptional FC, highlighting translationally regulated genes. D) Heatmap of differentially expressed genes (FC >1.5 and p-adjusted value <0.05) at the translational level.

**Fig. S8.**
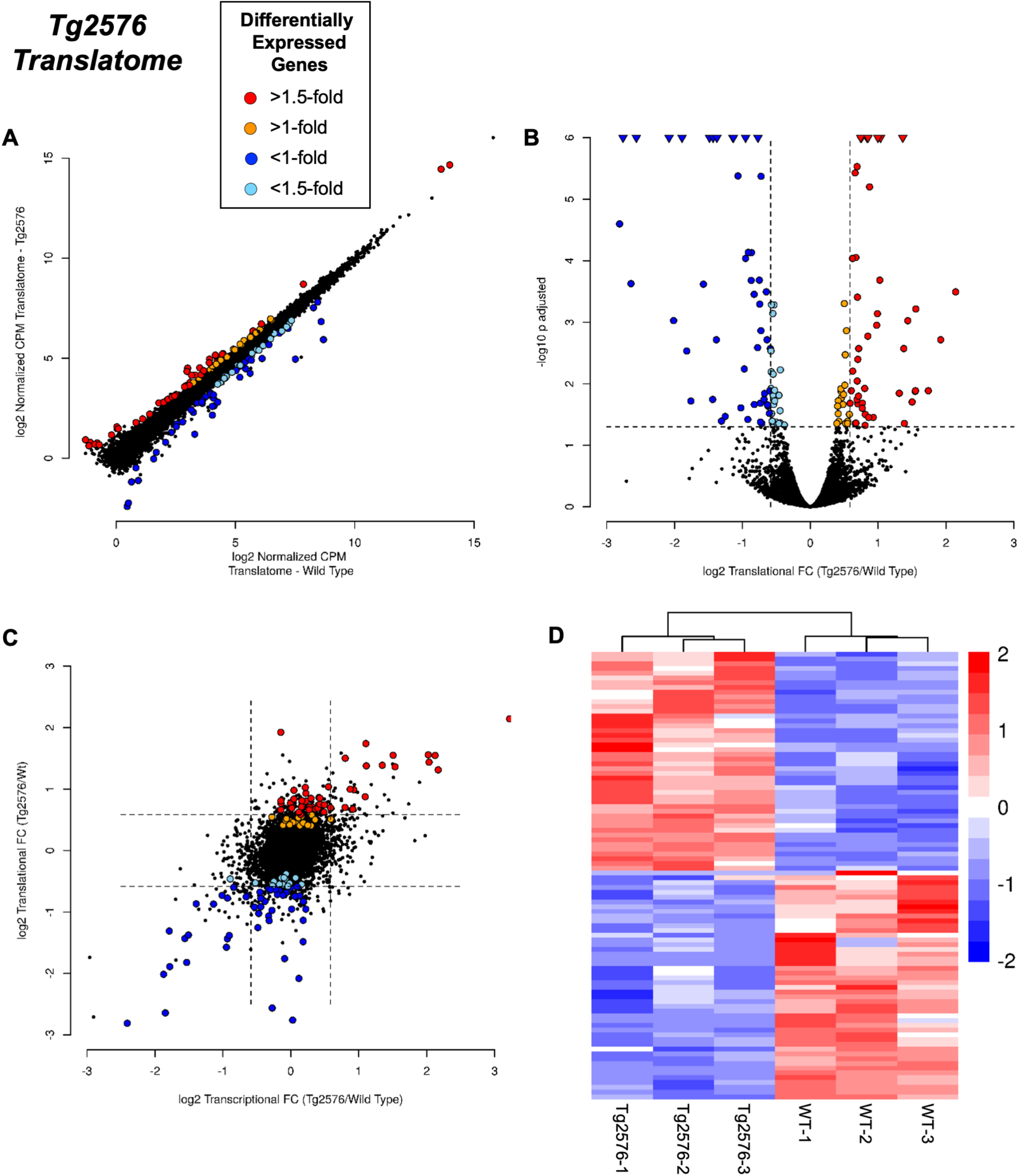
Differential expression analysis performed by *edgeR* for the translatome comparing Tg2576 mice versus their WT parental strain (B6;SJL). A) Scatter plot comparing normalized CPM expressions at the translational level between genotypes. B) Volcano plot of the relationship between FC and p-adjusted values. C) Scatter plot comparing translational vs transcriptional FC, highlighting translationally regulated genes. D) Heatmap of differentially expressed genes (FC >1.5 and p-adjusted value <0.05) at the translational level.

**Fig. S9.**
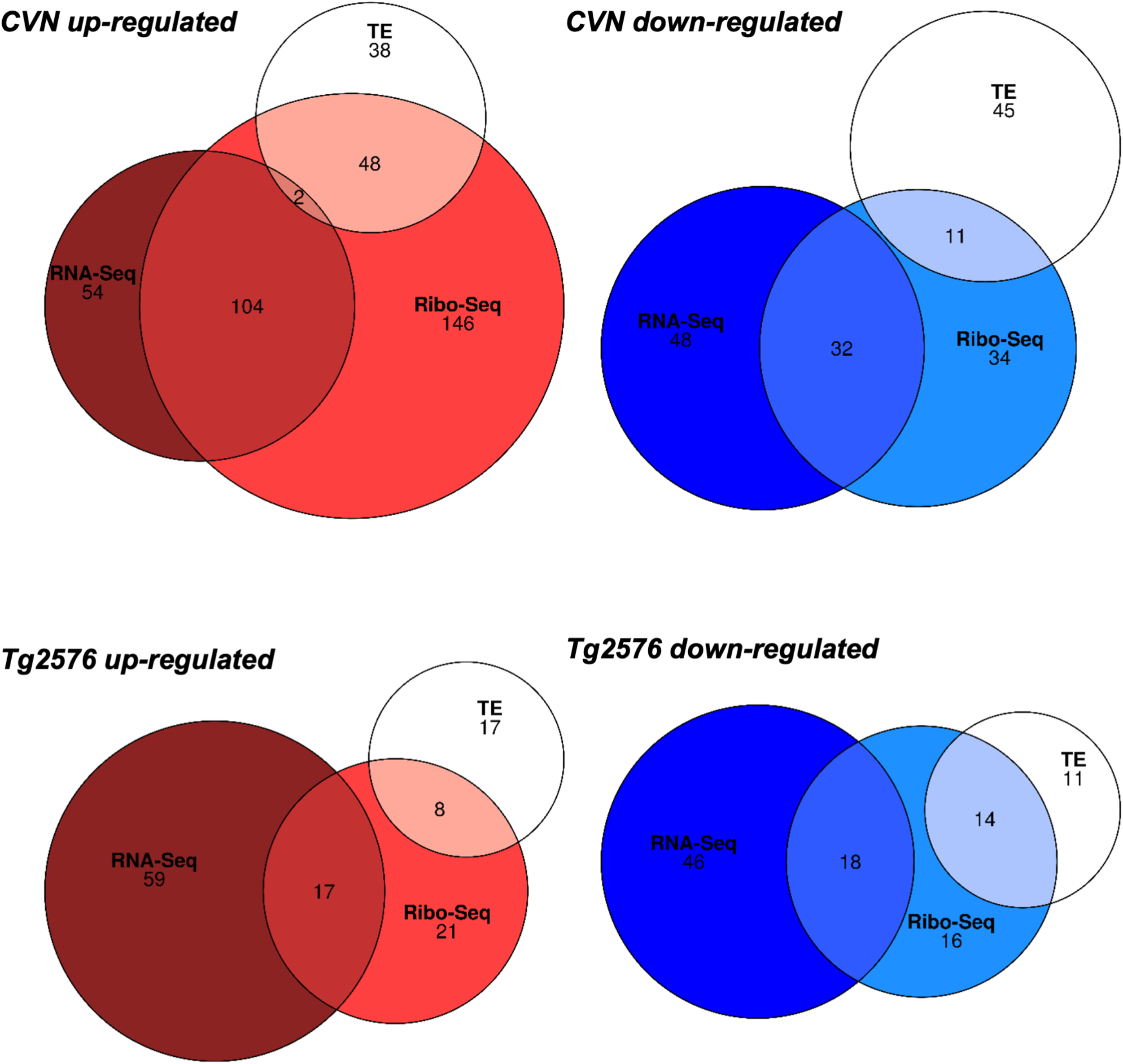
Intersection between differentially expressed genes at transcriptome (RNA-Seq), translatome (Ribo-Seq) and translational efficiency (TE) levels for CVN and Tg2576 mice.

**Fig. S10.**
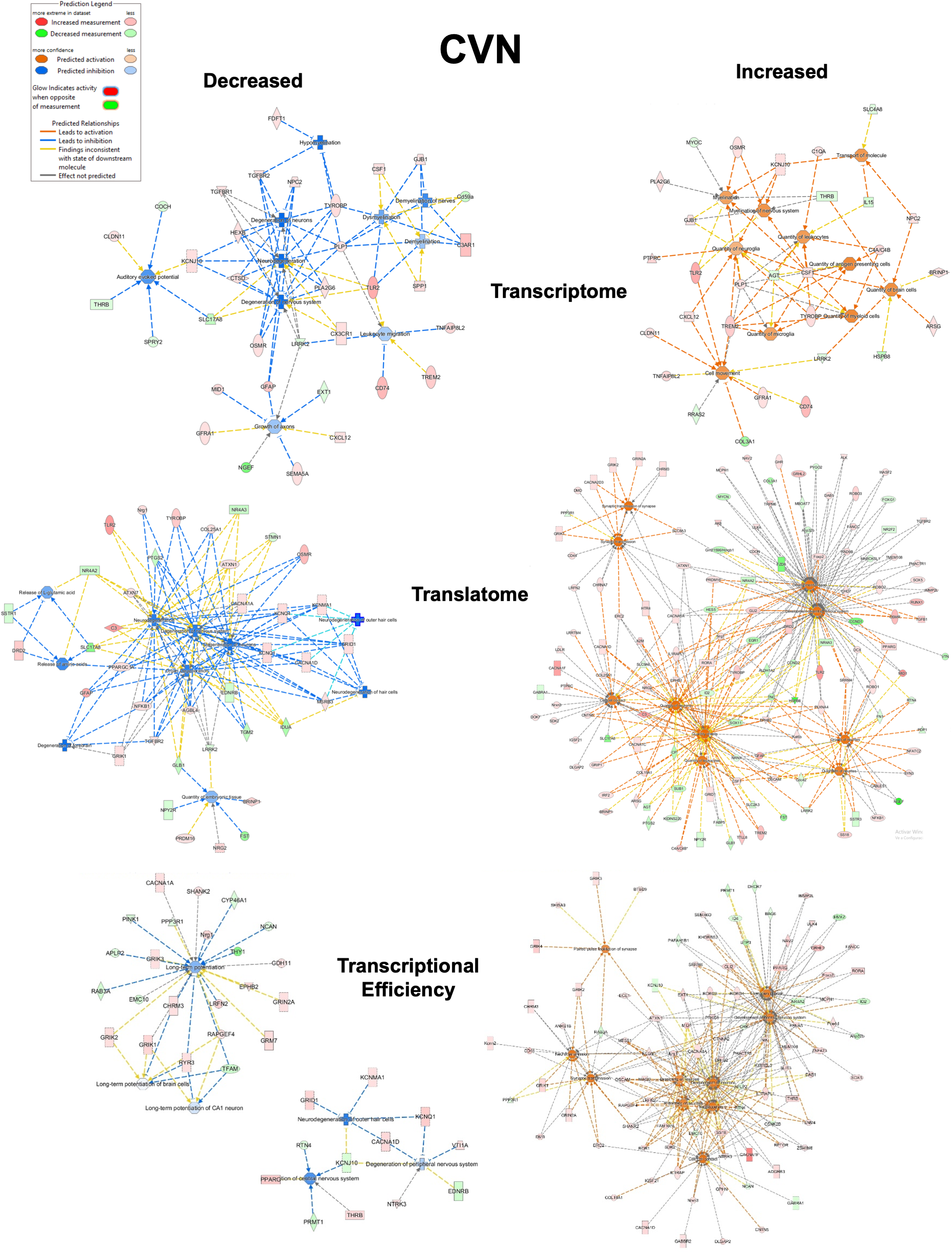
Modulated pathways in CVN mice. Regulated pathways showed in Figure 5 are reconstructed here showing affected genes and their relationship.

**Fig. S11.**
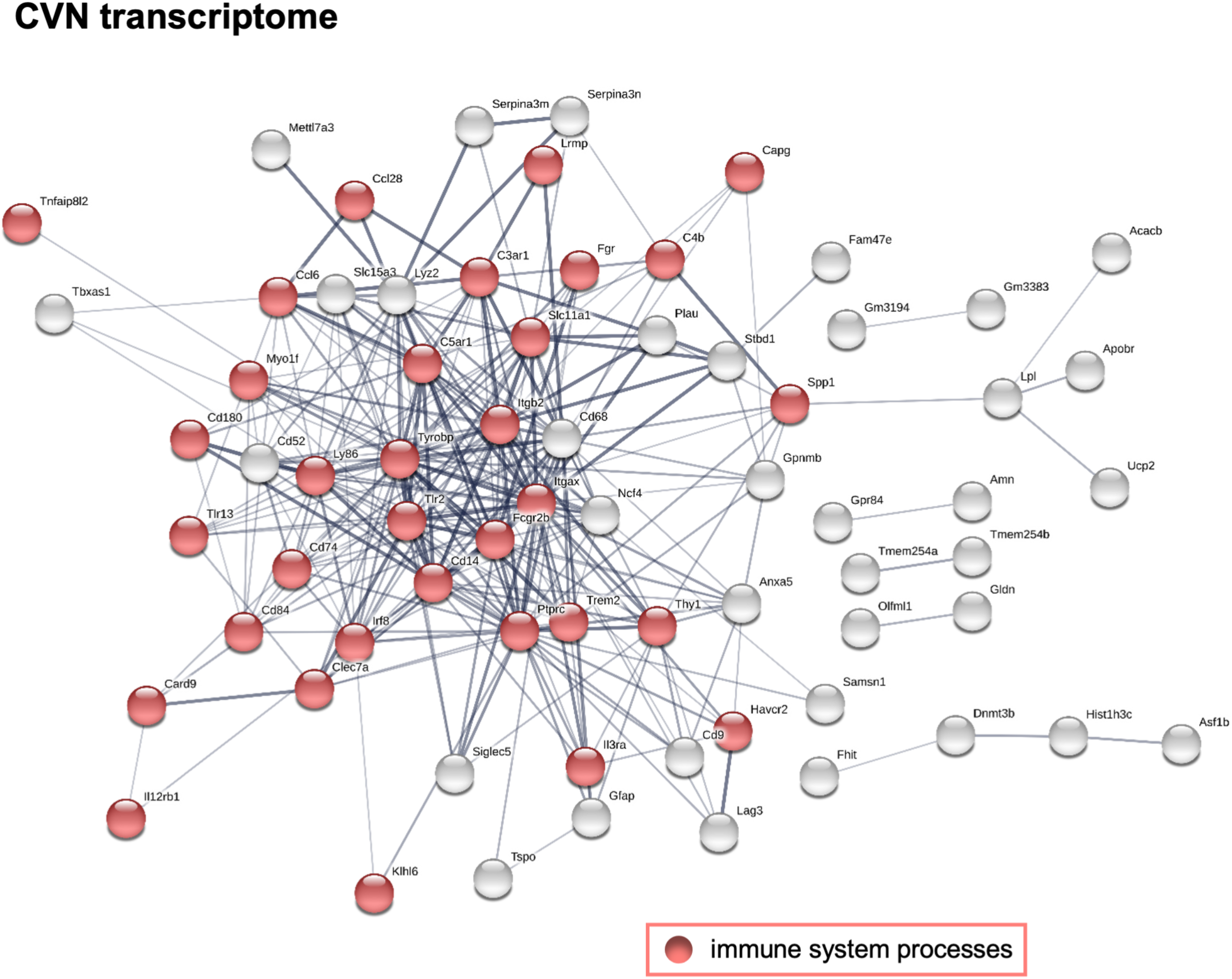
*STRING* network analysis for transcriptionally up-regulated genes (160 genes at FC >1.5 and p-adjusted value <0.05) in CVN mice. This network has significantly more interactions than expected (296 vs 53; p-adjusted value <1.0E-16). Red nodes are associated with immune system process (FDR = 2.67E-07). Disconnected nodes are hidden and line thickness indicates the strength of data support.

**Fig. S12.**
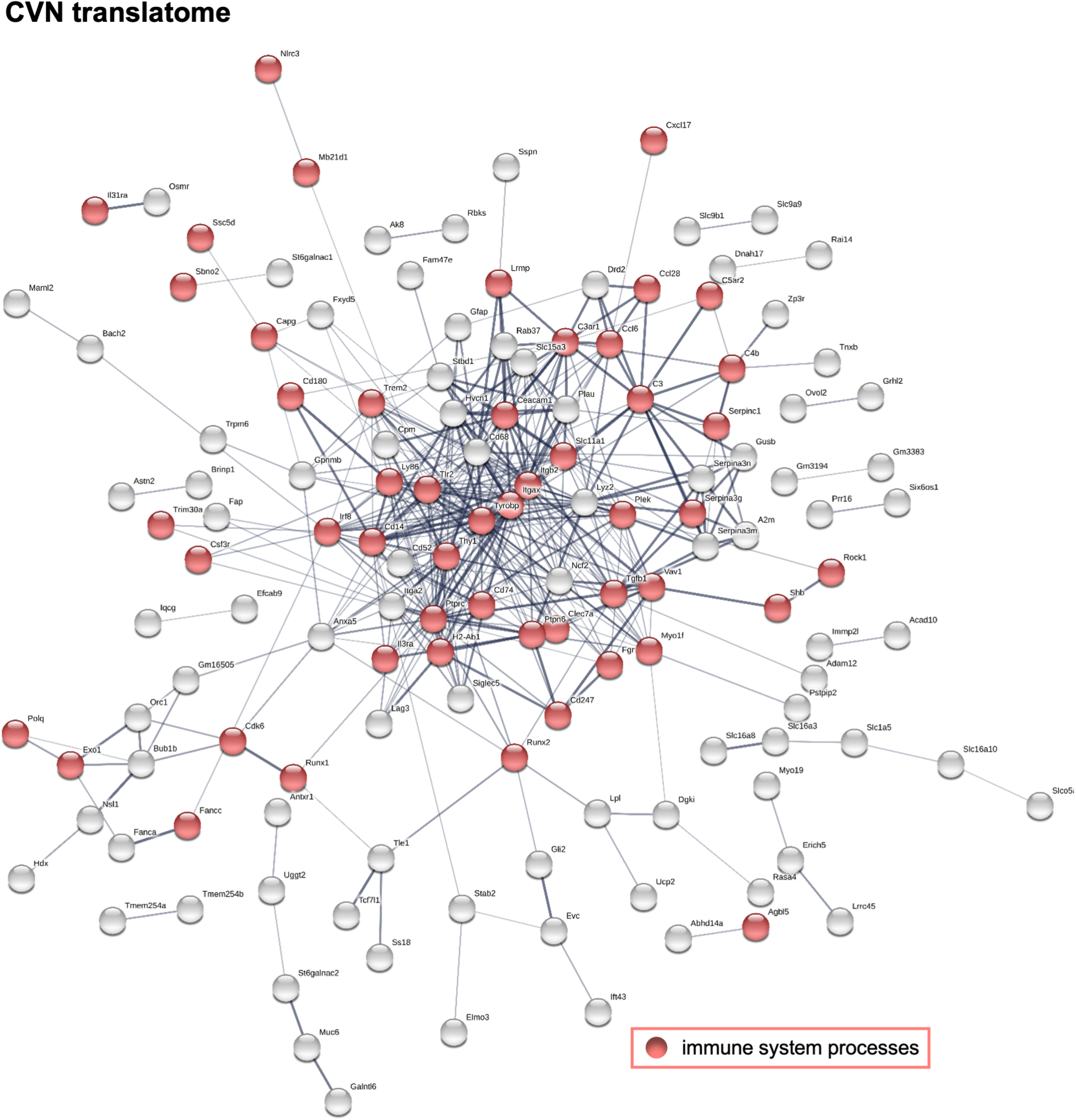
*STRING* network analysis for translationally up-regulated genes (300 genes at FC >1.5 and p-adjusted value <0.05) in CVN mice. Network has significantly more interactions than expected (431 vs 164; p <1.0E-16). Red nodes are associated with the immune system processes (FDR = 3.44E-07). Disconnected nodes are hidden and line thickness indicates the strength of data support.

**Fig. S13.**
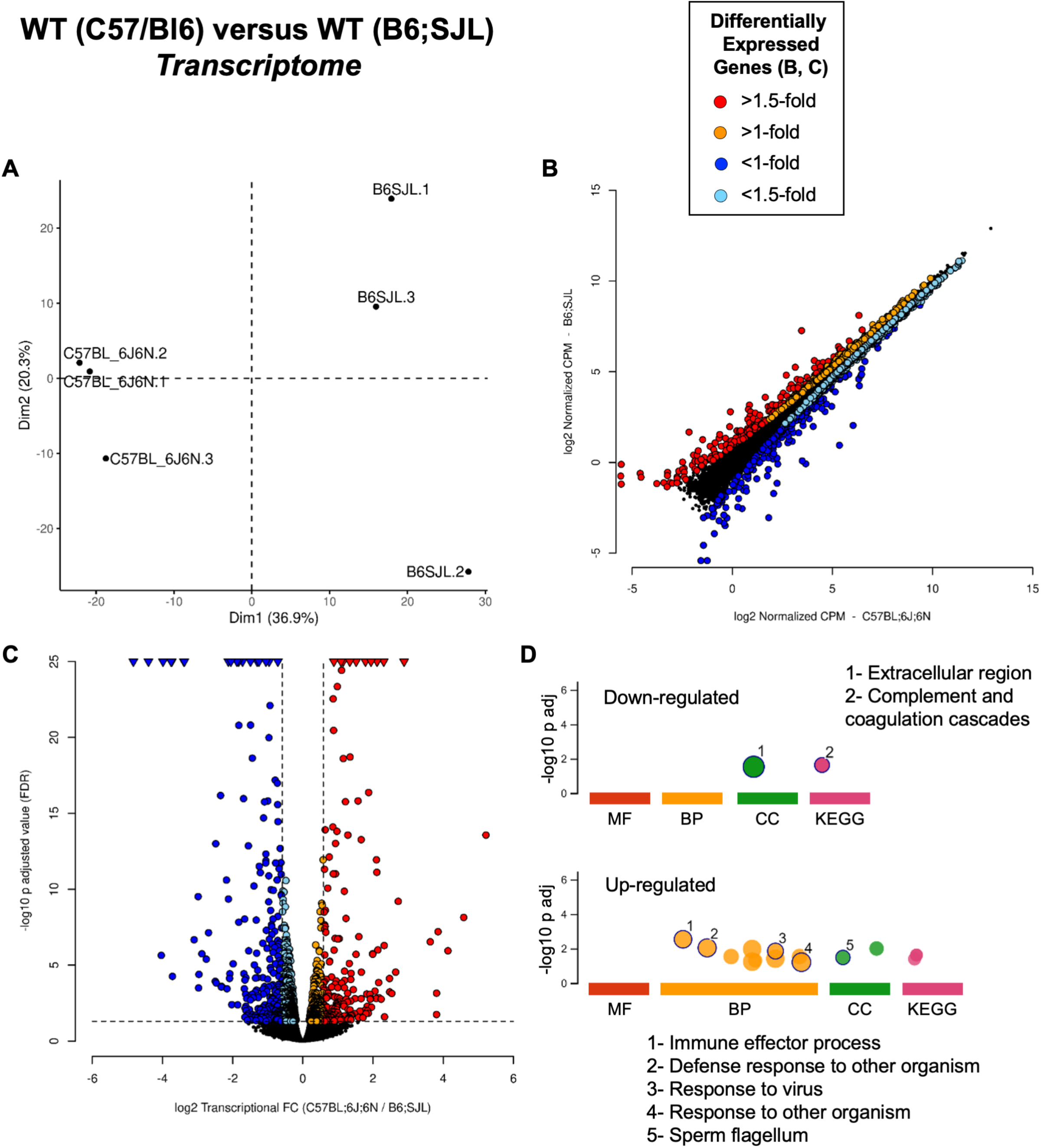
Differential expression analysis performed by *edgeR* for the transcriptome compartment comparing WT strains. A) Principal component analysis. B) Scatter plot comparing normalized CPM expressions at the transcriptome level between genotypes. C) Volcano plot showing the relationship between FC and p-adjusted values. D) Functional enrichment analysis of differentially expressed genes performed by *g:GOSt* from *g:Profiler*.

**Fig. S14.**
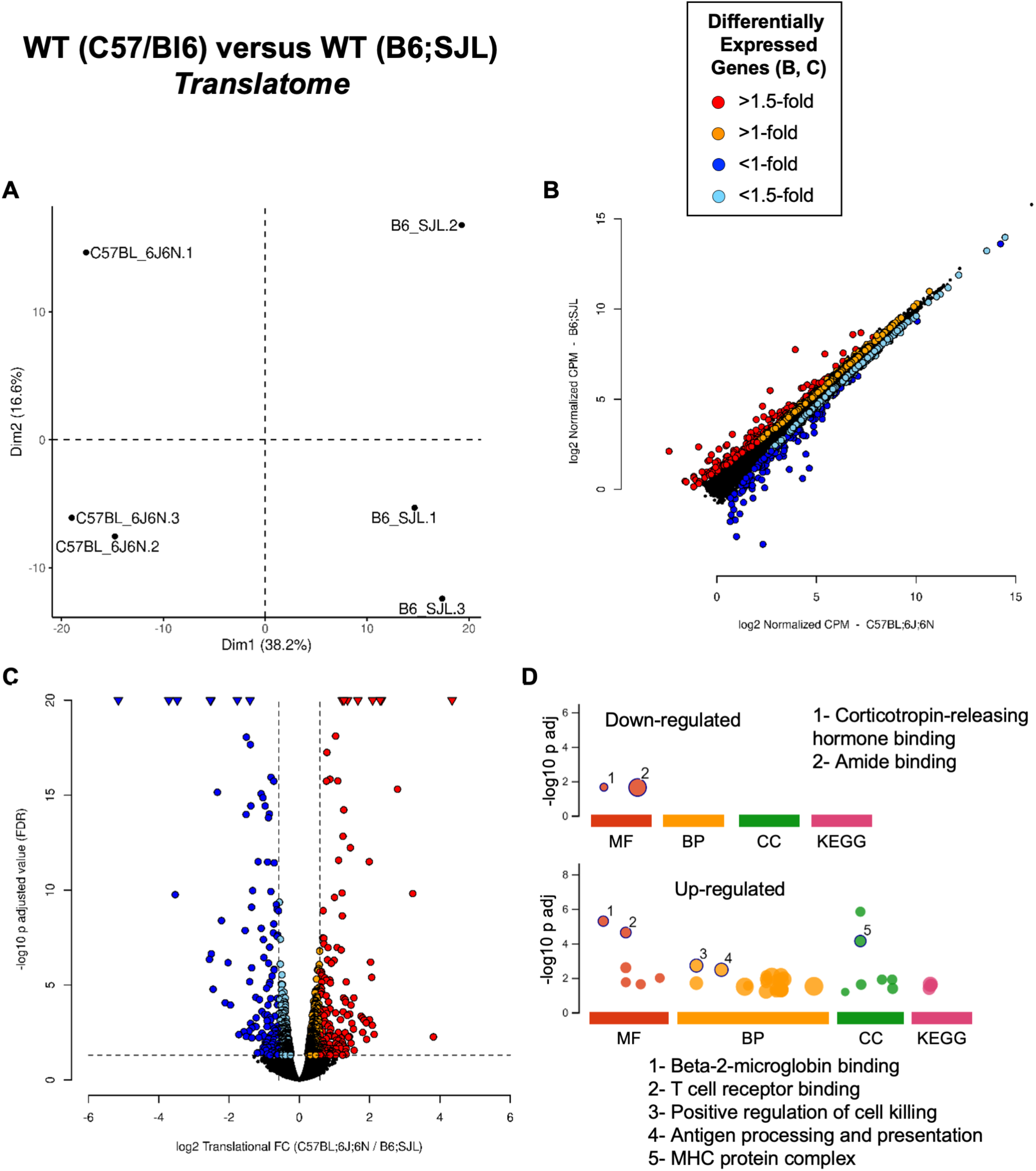
Differential expression analysis performed by *edgeR* for the translatome compartment comparing WT strains. A) Principal component analysis. B) Scatter plot comparing normalized CPM expressions at the translatome level between genotypes. C) Volcano plot showing the relationship between FC and p-adjusted values. D) Functional enrichment analysis of differentially expressed genes performed by *g:GOSt* from *g:Profiler*.

**Fig. S15.**
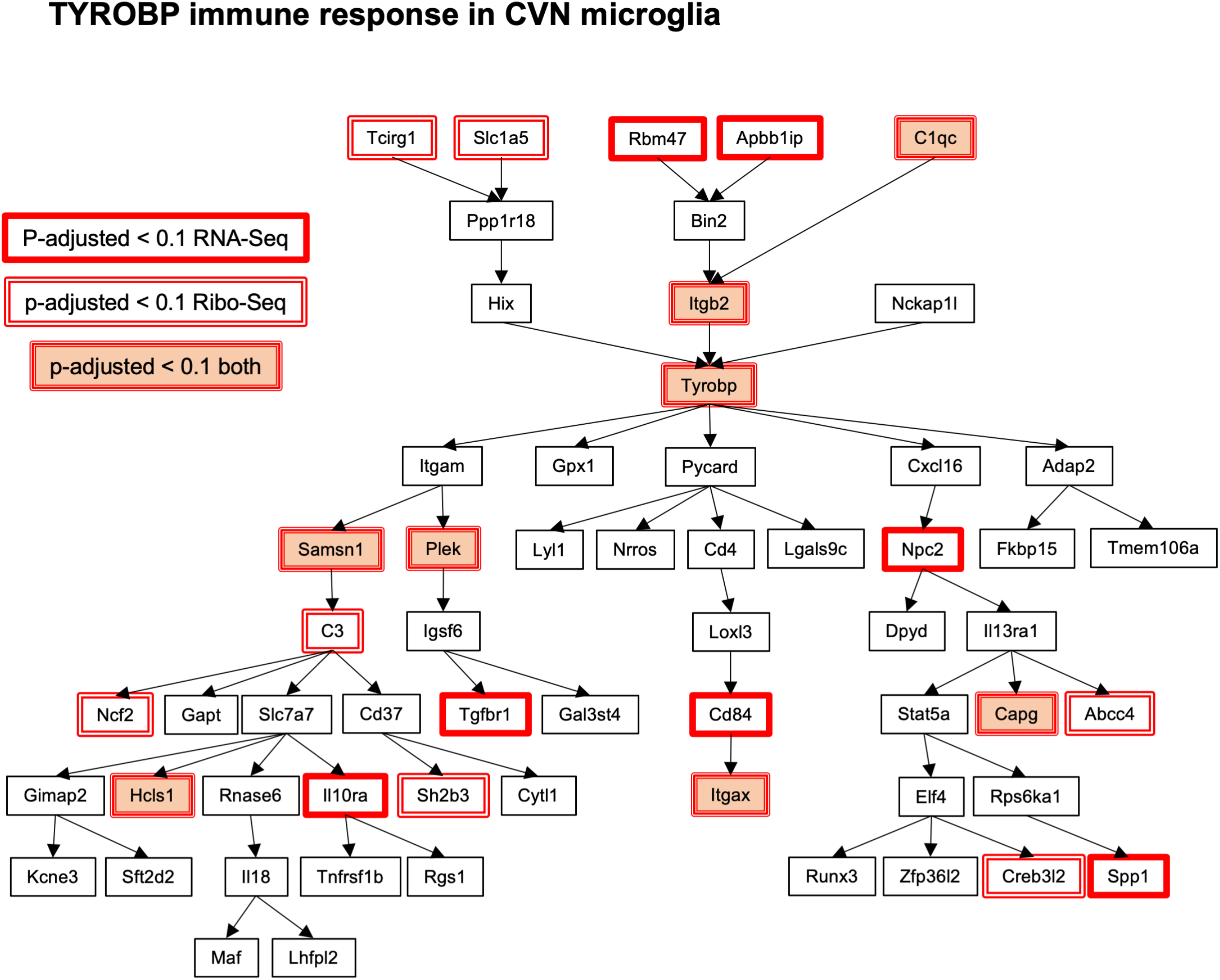
Gene expression regulation of TYRO protein kinase-binding protein (TYROBP) related immune response in CVN microglial cells. Pathway with direct and indirect causal inputs upstream and downstream of TYROBP was obtained from <u>WikiPathways</u>, reconstructed from Zhang B. et al. Cell (2013) and Ryu JK. et al. Nat Immunol (2018). Up-regulated genes at p-adjusted value <0.1 for transcriptome or translatome data sets are indicated, but no significant down-regulated genes were found.

